# Behavioral arousal organizes visual cortical population micro-states across the sleep-wake cycle

**DOI:** 10.64898/2026.09.29.755275

**Authors:** Jun-Ying Wang, Gerjan Huis in ‘t Veld, Kengo Takahashi, Kes Tijsseling, Mathis Bassler, Andreas Knoben, Cyriel M.A. Pennartz, Umberto Olcese

## Abstract

Brain states are conventionally defined as discrete categories, such as wakefulness, NREM sleep and REM sleep. Yet, whether these labels fully capture the organization of cortical population dynamics remains unclear. Using large-field-of-view two-photon calcium imaging of layer II/III pyramidal neurons in mouse primary visual cortex (V1), we show that population activity organizes into recurring micro-states that span conventional sleep-wake boundaries. A Hidden Markov Model fitted independently of behavioral state labels identified temporally stable micro-states with structured transition pathways. Micro-state organization was more strongly aligned with a continuous behavioral arousal axis, indexed by pupil diameter and orofacial motion, than with discrete NREM-wake categories, a relationship confirmed by mediation analysis. At the circuit level, two spatially intermingled but functionally opposing neuronal ensembles tracked this arousal axis in opposite directions. These findings establish behavioral arousal, rather than discrete sleep-wake state, as the primary organizing dimension of V1 population dynamics.

## Introduction

Patterns of neural activity greatly fluctuate across the sleep-wake cycle^1,2^, and brain-state dependent changes in brain activity profoundly influence cognitive processes including sensory processing, attention, and memory consolidation^3–8^. Brain states are conventionally classified via electrophysiological and behavioral signatures: slow oscillations, sleep spindles, and reduced behavioral responsiveness during non-rapid eye movement (NREM) sleep; theta oscillations and muscle atonia during rapid eye movement (REM) sleep; and desynchronized, low-amplitude activity during wakefulness^1,2,9,10^. Under this classical framework, wakefulness, NREM sleep, and REM sleep are conceived as discrete and globally expressed configurations of brain activity, with transitions between them usually portrayed as relatively abrupt and tightly regulated^11,12^. Historically, the co-expression of features conventionally associated with different sleep-wake states has been discussed primarily in the clinical context of parasomnias, which are often conceptualized as disorders of sleep-wake state dissociation^13^. However, accumulating evidence indicates that related dissociations can also occur under physiological conditions. For example, wake– and sleep-like dynamics can occur locally outside their globally defined state, for example in the form of sleep-like activity during wakefulness^14^, slow waves (typical of NREM) during REM sleep^15^ and distinct brain states simultaneously displayed by distant brain regions^16^. Beyond such cross-state dissociations, a single globally defined state can itself show heterogeneous neuronal dynamics. During wakefulness, cortical activity has long been known to fluctuate as a function of arousal and locomotion, with consequences for the ability to effectively process sensory stimuli^17,18^. Beyond wakefulness, even during NREM sleep, individual groups of neurons can show distinct, and even opposing, patterns of both activity and functional connectivity despite a common slow-wave signature at the population level^9,19–21^. Together, these findings suggest that conventional sleep-wake labels capture an important, yet only coarse-grained, dimension of brain-state dependent dynamics, and importantly fail to entirely specify the underlying, highly heterogeneous neuronal dynamics^22^. Consistent with this view, recent work suggests that fine-grained neuronal activity patterns can reveal a fine spatiotemporal micro-state organization that is obscured by conventional global state scoring^23^. However, how such micro-states are embedded into the wake-sleep cycle, what characterizes them in terms of patterns of neuronal activity and what governs the transition between micro-states remain open questions.

Here we address this by characterizing neuronal population activity patterns across spontaneous sleep-wake fluctuations in primary visual cortex (V1). Using large-field-of-view two-photon calcium imaging, we simultaneously recorded hundreds of layer II/III pyramidal neurons during spontaneous sleep-wake fluctuations in head-fixed mice, while monitoring pupil diameter and orofacial movements as continuous indices of behavioral arousal. We then used a Hidden Markov Model (HMM), fitted solely to neuronal activity and without incorporating conventional sleep-wake labels, to identify recurring population micro-states. These micro-states were expressed both within and across NREM sleep and wakefulness, indicating that global sleep-wake labels did not uniquely specify V1 population activity. Instead, the organization of micro-states was more coherently captured by a continuous behavioral arousal axis derived from pupil diameter and orofacial motion. At the circuit level, this axis was represented by two opposing types of neuronal ensembles, whose activity increased or decreased progressively with arousal. Together, these findings reveal that, in V1, brain states are determined by a repertoire of recurring population micro-states whose organization is more continuously structured by behavioral arousal than by discrete global sleep-wake state labels.

## Results

### A pupil-based classifier reliably distinguishes NREM sleep from Wakefulness

To investigate the relationship between cortical population dynamics and brain state, we recorded from head-fixed mice across two complementary datasets (**Fig. 1A**). In Dataset A (n = 7 sessions from 7 mice), we used large field-of-view (FOV) two-photon calcium imaging to monitor GCaMP6f-expressing layer II/III IT pyramidal neurons in primary visual cortex (V1) (**Fig. 1B**, **Fig. S1**). This approach provides dense sampling of a local neuronal population at single-cell resolution, making it well-suited to characterizing coordinated population dynamics across brain states. Mice were head-fixed and imaged during spontaneous behavior in complete darkness with no visual stimulation, allowing natural alternation between wakefulness and sleep within a single ∼2-hour session (see **Methods**). Pupil diameter and orofacial motion energy were recorded using an infrared face camera and quantified with FaceMap^24^ (**Fig. 1C**). Neural data was visualized using Rastermap^25^ and aligned to co-varying fluctuations in both pupil diameter and orofacial motion (**Fig. 1D**). Fluctuations in pupil size have emerged as a reliable proxy for brain state identification^26,27^; however, electrophysiological recordings remain the gold standard for brain state determination^28^, and Dataset A lacked such a measure. Therefore, to validate the pupil-based estimation of brain state, we assembled Dataset B (n = 6 sessions from 2 mice), in which Neuropixels probes were implanted in V1 to provide simultaneous LFP recordings alongside the same face camera setup (**Fig. 1A**). Manual scoring of LFP traces identified NREM sleep or Wakefulness (WAKE) epochs (**Fig. 1E**). Putative REM-like epochs were infrequent and could not be unambiguously classified without concurrent electromyography (EMG); they were therefore labelled Unclassified and excluded, restricting subsequent analyses to NREM and WAKE (see **Methods**). Pupil diameter distributions differed markedly between these LFP-classified states (**Fig. 1F–G**): NREM epochs showed a broad, flat distribution of normalized pupil diameter, reflecting large fluctuations, whereas WAKE epochs were characterized by a narrow distribution shifted towards larger values, consistent with sustained pupil dilation during wakefulness.

**Figure 1.**
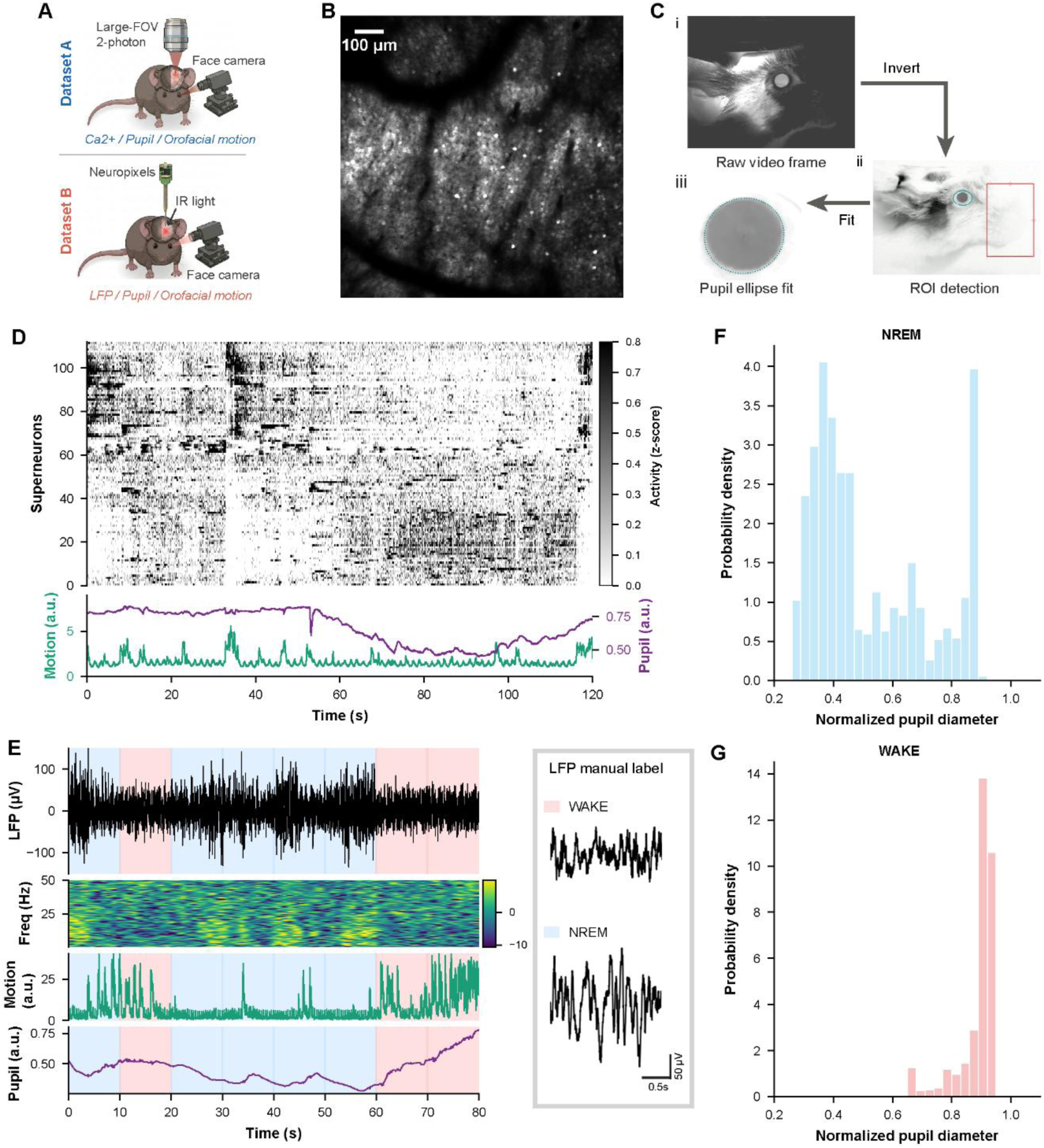
Experimental design and representative recordings. **(A)** Schematic of the two recording configurations. Dataset A: head-fixed mice were imaged with a large field-of-view (FOV) two-photon microscope offering single-cell resolution, while an infrared face camera simultaneously captured pupil diameter and orofacial motion (Ca²⁺ / Pupil / Orofacial motion; n = 7 sessions from 7 mice). Dataset B: a separate cohort of head-fixed mice was implanted with a Neuropixels probe and recorded under infrared (IR) illumination with the same face-camera setup (LFP / Pupil / Orofacial motion; n = 6 sessions from 2 mice). **(B)** Representative two-photon FOV from Dataset A showing GCaMP6f-expressing layer II/III IT pyramidal neurons in primary visual cortex. Scale bar, 100 µm. **(C)** Pupil and orofacial motion analysis pipeline using FaceMap. (i) Raw infrared video frame. (ii) Frame inversion and definition of two ROIs: an eye ROI and an orofacial ROI for motion energy computation. (iii) Ellipse fitting to the eye ROI to extract instantaneous pupil area. **(D)** Representative Dataset A session (obtained by 2-photon imaging). Top: superneuron activity heatmap (z-scored ΔF/F) sorted by Rastermap; each row represents a superneuron (bin size = 5 neurons). Bottom: simultaneously recorded normalized orofacial motion (green) and normalized pupil diameter (purple). **(E)** Representative Dataset B session. From top to bottom: average LFP trace recorded from visual cortex (µV); LFP spectrogram (relative power, dB); normalized orofacial motion (green); normalized pupil diameter (purple). Background shading indicates manually assigned LFP-based brain state labels (blue: NREM; red: WAKE). Right insets: representative zoom-in on LFP traces extracted from a WAKE epoch (top) and a NREM epoch (bottom), illustrating the characteristic desynchronized and slow-oscillatory waveform patterns, respectively. Scale bar, 50 µV / 0.5 s. **(F, G)** Pupil diameter distributions during representative NREM (**F**) and WAKE (**G**) epochs from the same Dataset B session. Histograms show the probability density of normalized pupil diameter within each epoch (blue: NREM; red: WAKE; total histogram area (bin width X bin height) = 1).

While pupil diameter distributions differed between brain states at the single-epoch level, plotting all LFP-labeled epochs according to their median pupil diameter and pupil-diameter distribution width revealed substantial overlap between NREM and WAKE, precluding reliable classification based on these two features alone (**Fig. S2A**). To improve discriminability, we trained a logistic regression classifier on Dataset B, using four pupil-derived features as predictors and LFP-derived brain state labels as ground truth (**Fig. 2A**). The feature set comprised four pupil-derived measures capturing both the static level and temporal dynamics of pupil size within a 60-second epoch (see **Methods**), all of which meaningfully contributed to the classifier (**Fig. S2C**).

**Figure 2.**
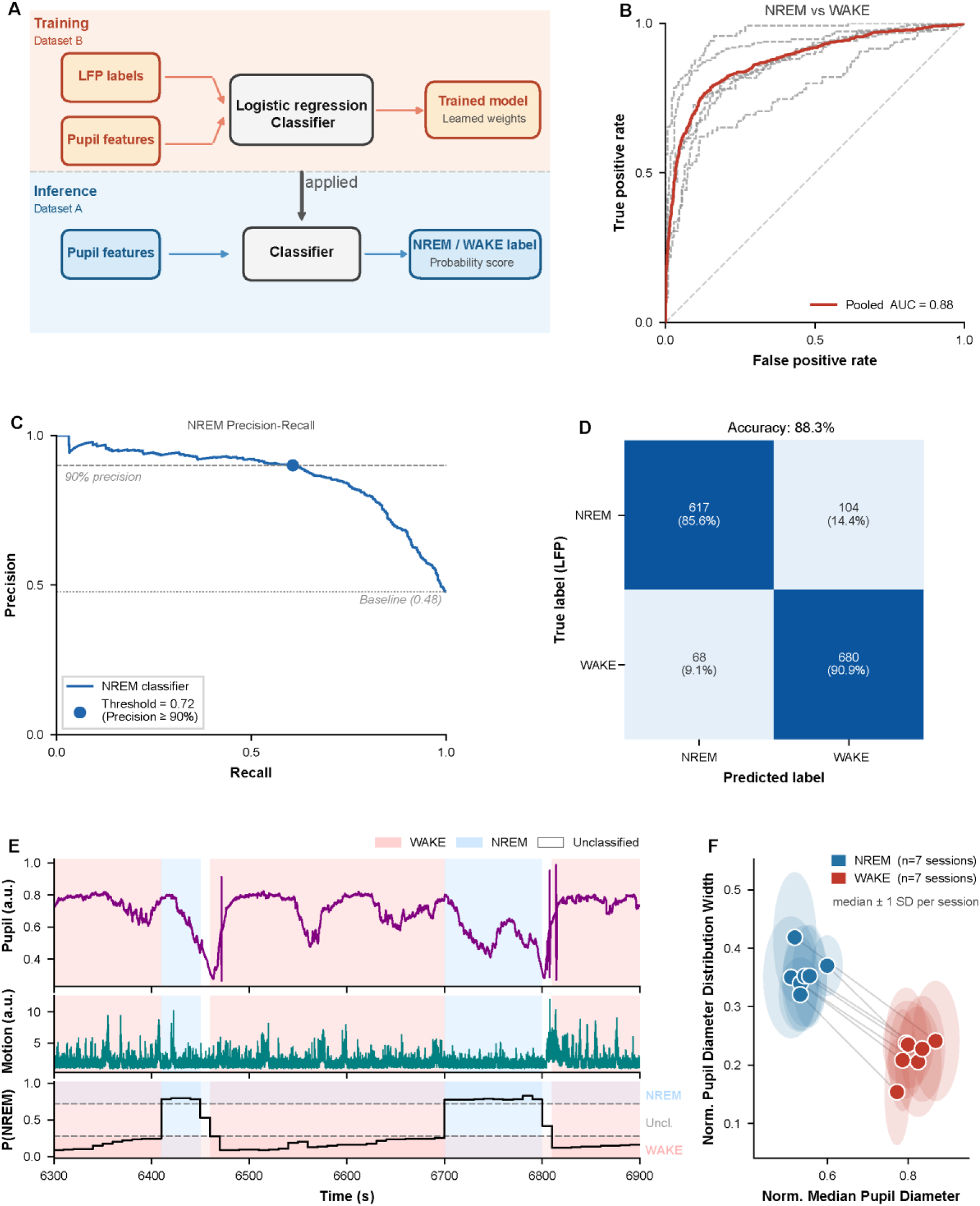
Logistic regression classifier for pupil-based brain state inference. **(A)** Schematic of the classification pipeline. A logistic regression classifier was trained on LFP-labelled pupil features from Dataset B (top) and applied to Dataset A for inference (bottom). **(B)** Receiver operating characteristic (ROC) curves for NREM vs WAKE classification. Grey dashed lines show per-session curves (n=6); the solid red line shows the pooled curve. Mean per-session AUC = 0.88 ± 0.06; pooled AUC = 0.88. AUC, area under the curve**. (C)** Precision-recall curve for the NREM classification using pooled Dataset B predictions. Each point represents classifier performance at a different posterior-probability threshold for assigning an epoch to NREM. The filled circle indicates the selected threshold, P(NREM) = 0.72, the lowest threshold at which NREM precision was at least 90%. The dashed horizontal line indicates the 90% precision; the dotted line shows the no-skill baseline, equal to NREM prevalence. **(D)** Confusion matrix for the logistic regression classifier evaluated on Dataset B, pooled across all sessions, after applying the posterior probability threshold (≥ 0.72). Values show epoch counts and row-normalized percentages. **(E)** Representative time series from a Dataset A session showing normalized pupil diameter (top), normalized orofacial motion energy (middle), and P(NREM) output from the logistic regression classifier (bottom). Transient sharp deflections in the pupil-diameter trace reflect brief eyelid occlusions during blinks, which were retained during feature extraction (see **Methods**). Background shading indicates the predicted state (blue: NREM; red: WAKE; blank: Unclassified) for each 60-second epoch. Dashed lines mark the classification thresholds (P(NREM) = 0.72 and 0.28). Unclassified windows (no shading) fall below the confidence threshold. **(F)** Cross-session summary of LR-predicted brain states in Dataset A. Each point represents the per-session centroid for NREM (blue) and WAKE (red) in the pupil feature space; ellipses show ±1 SD of epoch-level features within each session. Lines connect NREM and WAKE centroids from the same session.

Leave-one-session-out cross-validation on Dataset B confirmed the ability of the classifier to reliably discriminate between NREM and WAKE, yielding a pooled area under the curve (AUC) of 0.88 ± 0.06 (**Fig. 2B**). To ensure high-confidence state assignments, we applied a posterior probability threshold of 0.72, corresponding to the point on the precision–recall curve at which NREM precision reached ≥ 90% (**Fig. 2C**, see **Methods**). Restricting Dataset B predictions to this high-confidence subset yielded an overall classification accuracy of 88.3% (NREM: 85.6%; WAKE: 90.9%; **Fig. 2D**). Accordingly, in the same representative session as Fig. S2A, now colored by classifier-predicted label, this thresholding procedure markedly sharpened the boundary between NREM and WAKE in the pupil feature space (**Fig. S2B**).

We then applied the trained model to Dataset A (n=7 sessions), for which no LFP recordings were available. The NREM probability P(NREM) time series captured state transitions consistent with the canonical signatures of NREM and Wakefulness (**Fig. 2E**): NREM epochs were characterized by lower pupil diameter with high variability and reduced orofacial motion, whereas WAKE epochs showed elevated pupil diameter and increased orofacial motion energy. Notably, orofacial motion was not included as a classifier feature, and thus provides an independent behavioral validation of the predicted brain states. At the single-session level, NREM and WAKE epochs remained separable in the pupil feature space after applying the classifier (**Fig. S2D**). This separation was consistent across sessions, with WAKE centroids shifted towards higher median diameter and lower distribution width relative to NREM (**Fig. 2F**).

### V1 population activity organizes into temporally stable and behaviorally distinct micro-states

Having identified NREM and WAKE epochs, we first visually inspected V1 population activity patterns within these epochs using Rastermap^25^ (**Fig. 3A**). At the population level, heterogeneous patterns could be observed within both WAKE and NREM. For example, within NREM some epochs showed broad activations across a large proportion of neurons (**Fig. 3Ai**), while others were characterized by sparse, transient activations involving a small fraction of neurons (**Fig. 3Aii**). A comparable degree of heterogeneity was observed in WAKE (**Fig. 3Aiii, iv**). While WAKE epochs lacked the long periods of coordinated and sustained activity that could be observed in NREM (cf. **Fig. 3Ai, iii**), we observed instances in which population activity patterns appeared to be similar between WAKE and NREM epochs (**Fig. 3Aii, iv**). Together, these observations suggest that macro-state identity alone is insufficient to capture the full diversity of V1 population dynamics. This motivated us to investigate whether recurring micro-states could be identified within wakefulness and sleep, and whether such micro-states were distinct or shared between macro-states. To this end, we applied a Hidden Markov Model (HMM) to the ΔF/F population activity to identify stable patterns of activity independently from the assigned macro-state.

**Figure 3.**
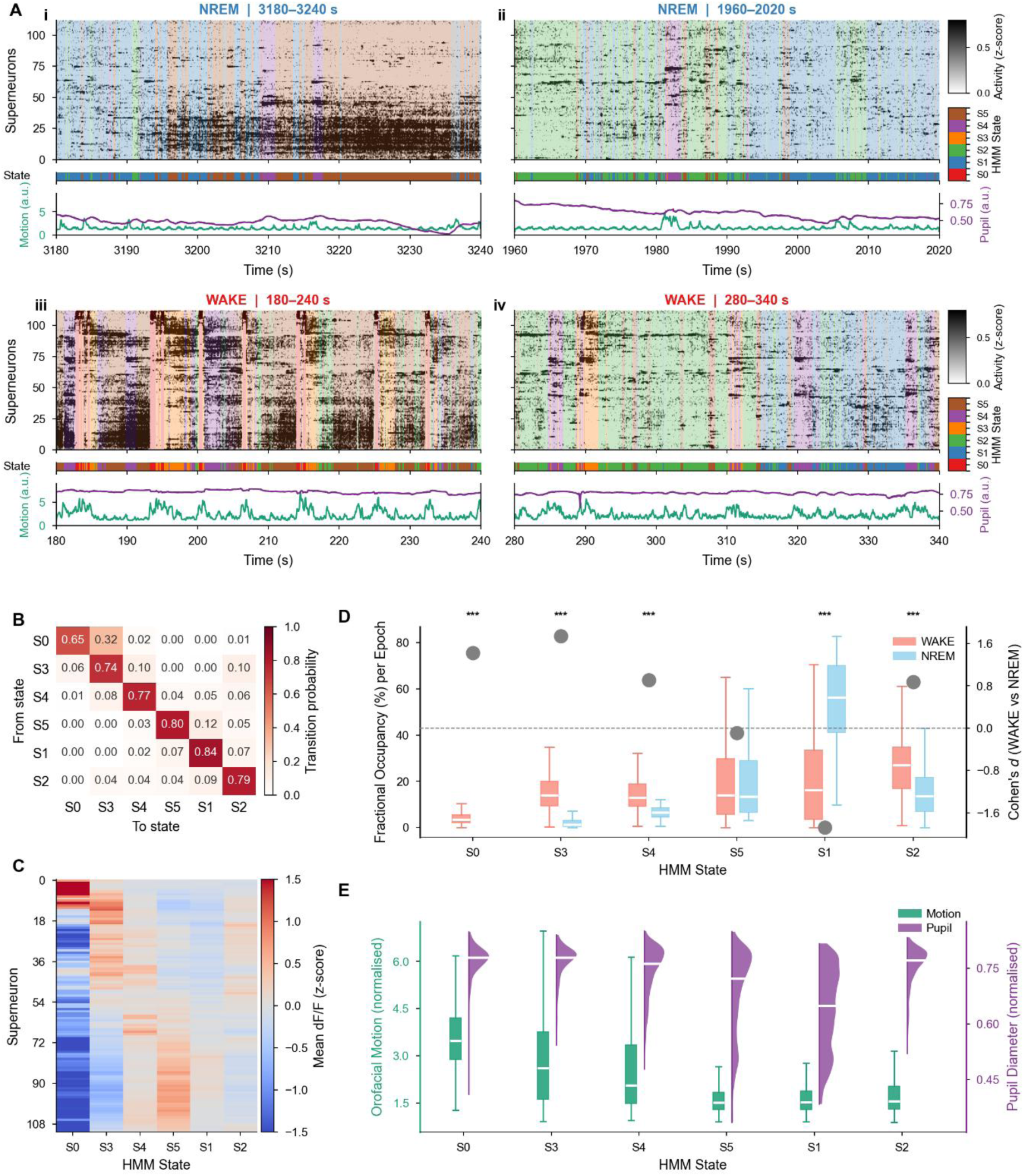
V1 population activity organizes into recurring micro-states with distinct temporal, structural, and behavioral signatures. **(A)** Representative V1 population activity for four 60-second epochs: two NREM epochs (i, ii) and two WAKE epochs (iii, iv). Each panel shows population activity (Rastermap, top; gray-scale, z-scored ΔF/F), HMM state sequence (middle bar), and simultaneously recorded orofacial motion (green) and normalized pupil diameter (purple; bottom). Colored overlays indicate HMM micro-state identity at each timepoint. **(B)** Transition probability matrix for a representative session (K = 6 states), with states ordered by transition similarity. **(C)** Neural activity fingerprints of each HMM state (states ordered as in B). Heatmap shows mean z-scored ΔF/F for individual neurons grouped into superneuron bins for visualization (rows, sorted by Rastermap embedding order) for each state (columns). **(D)** Fractional occupancy (% per 60-second epoch) of each HMM state during WAKE (salmon) and NREM (blue) epochs (states ordered as in B). Asterisks indicate significant WAKE vs. NREM differences (Mann-Whitney U test, FDR-corrected; *p < 0.05, **p < 0.01, ***p < 0.001). Grey dots show Cohen’s *d* effect size (WAKE vs. NREM) for each state (right axis; dashed line = 0), with positive values indicating greater WAKE occupancy. **(E)** Behavioral signatures of HMM micro-states (states ordered as in B). For each state, box plots show the distribution of normalized orofacial motion (green, left axis; box: median and IQR; whiskers: 1.5× IQR) and half-violin plots show the distribution of normalized pupil diameter (purple, right axis). Group differences were assessed by Kruskal–Wallis test (motion: H = 9641.9, ε² = 0.16; pupil: H = 13374.9, ε² = 0.22; both p < 0.001). Post-hoc Dunn’s tests with Bonferroni correction revealed significant pairwise differences in 15/15 state pairs for motion and 14/15 state pairs for pupil diameter (all except S0 vs. S3).

For each session, the optimal number of micro-states (K) was determined using a multi-stage selection procedure (**Fig. S3A-F; Methods**). First, biological and model-quality criteria were applied to define a per-session candidate pool of K values. Because the candidate pools were contiguous ranges beginning at the minimum admissible K, we used the upper bound of each pool (K_up_) as the session-level candidate number of micro-states for cross-session optimization. This retained the most complex model supported by the within-session biological and model-quality criteria while avoiding further expansion of the cross-session search space. To ensure that the identified micro-states captured patterns of activity that generalized across animals rather than reflecting session-specific features, we selected the combination of K_up_ values across all seven sessions that maximized cross-session reproducibility, quantified as the mean pairwise Pearson correlation of state feature profiles – with each micro-state characterized by its fractional occupancy, self-transition probability, activity entropy, and population synchrony – matched across sessions using the Hungarian algorithm^29^. Because HMMs were fitted independently per session, the optimal K differed across sessions (final K range: 5–8 states). This combination yielded a mean pairwise cross-session correlation of 0.80, indicating highly consistent micro-state structure across animals (**Fig. S3G-I**; **Methods**).

Having established cross-session consistency among micro-states, we next characterized the structure of the identified micro-states. Within a representative session (K = 6), self-transition probabilities ranged from 0.65 to 0.84 (**Fig. 3B**); across all sessions, self-transition probabilities ranged from 0.40 to 0.85 (mean ± SD: 0.70 ± 0.13), indicating that micro-states were generally maintained across consecutive frames, with variability in temporal stability reflecting state-specific differences in neural dynamics. Micro-state durations were right-skewed across all states and sessions (skewness range 1.30–15.39, median duration 0.21 s, mean duration 0.51 s), consistent with brief, frequent state visits punctuated by occasional sustained episodes. Examining the off-diagonal structure of the transition matrix revealed that non-self-transitions were likewise non-random: specific state pairs showed highly frequent transitions, while others showed near-zero transition probability. In the representative session (**Fig. 3B**), micro-state S0 almost exclusively transitioned to S3 (probability: 0.92), and micro-state S5 showed a strong preference for transitioning to S1 (probability: 0.59), while S1 and S2 showed frequent reciprocal transitions. This structured transition topology was largely conserved across WAKE and NREM epochs, suggesting that preferred transition pathways reflect intrinsic properties of V1 population dynamics (**Fig. S4**). Macro-state identity nonetheless modulated the relative weighting of specific transitions within this stable topology. For example, the S5→S1 transition was stronger during NREM than WAKE (**Fig. S4D**), indicating that macro states shape the expression of micro-state dynamics without fundamentally reorganizing their structure. Complementing this temporal organization, each micro-state was characterized by a distinct population-level activity fingerprint, with specific subsets of neurons consistently elevated or suppressed relative to the session mean (**Fig. 3C**). Across all sessions, mean activity patterns were consistently similar across repeated occurrences of the same micro-state (within-state cosine distance: 0.79 ± 0.05), and different between micro-states (between-state cosine distance: 1.17 ± 0.04), yielding a separation ratio of 1.48 ± 0.11 (with a ratio > 1 indicating between-state distances exceed within-state distances; see **Methods**). Together, the temporal stability, structured transitions, and stereotypical neural signatures confirm that HMM-identified micro-states reflect genuine recurring population dynamics.

To assess whether micro-states mapped onto conventional macro-states (WAKE and NREM), we quantified the fractional occupancy of each micro-state within NREM and WAKE epochs, defined as the proportion of frames assigned to that micro-state relative to the total frames within that macro-state (**Fig. 3D**). No micro-state was entirely absent from either macro-state, though fractional occupancy (FO) was highly heterogeneous: for example, in the representative session shown earlier micro-state S0 displayed near-exclusive association with WAKE epochs (FO_WAKE_: 3.67%, FO_NREM_: 0.30%), while micro-state S1 showed the strongest sleep preference (FO_WAKE_: 19.99%, FO_NREM_: 53.77%). We quantified each micro-state’s WAKE or NREM preference across sessions using Cohen’s *d* for the difference in fractional occupancy between WAKE and NREM^30^. Positive values of the Cohen’s *d* indicate WAKE preference and negative values indicate NREM preference **(Fig. 3D, right axis)**. Across all recorded sessions, micro-states showed a broad distribution in Cohen’s *d* values (**Fig. S6A**), indicating a graded rather than categorical relationship between micro-states and macro-state identity (range: −2.36 to +1.74; mean |*d*| = 0.85 ± 0.51). Beyond their macro-state preference, micro-states were further distinguished by the co-occurring level of pupil dilation and orofacial motion energy. In the representative session, all micro-states differed significantly in both orofacial motion energy and pupil diameter (Kruskal-Wallis test: motion, H = 9641.9, ε² = 0.16; pupil, H = 13374.9, ε² = 0.22; both p < 0.001; **Fig. 3E**). This pattern was replicated across sessions: both measures varied significantly across micro-states in every session tested (all Kruskal-Wallis test, p < 0.05; median ε² = 0.155 for motion, IQR = 0.087; median ε² = 0.153 for pupil diameter, IQR = 0.041). Together, these findings indicate that micro-states were associated with distinct, although partially overlapping, behavioral signatures.

### Neural micro-states are organized along a behavioral arousal axis

The micro-states that we identified in population activity showed a mixed, non-trivial relationship with behavioral macro-states, pupil-based arousal and orofacial motion. To further explore the relationship between micro-states and behavioral features, we first plotted the median normalized pupil diameter and median normalized orofacial motion of each micro-state (n = 48 states, 7 sessions) (**Fig. 4A**). Separately for each session, pupil and motion medians were z-scored across micro-states to remove between-session differences in baseline pupil size and movement level before pooling across animals. Micro-states were organized into five reproducible clusters in this two-dimensional pupil-motion (PM) space, identified by k-means clustering (5 clusters; silhouette score = 0.57), with the number of clusters selected as the solution that best balanced statistical compactness and behavioral interpretability (**Fig. S5A-B**): a high-motion, high-pupil cluster (HM-HP); an intermediate-motion, high-pupil cluster (IM-HP); a low-motion, high-pupil cluster (LM-HP); a low-motion, intermediate-pupil cluster (LM-IP); and a low-motion, low-pupil cluster (LM-LP). The cluster structure was consistent across sessions and robust to the number of HMM states used per session, as confirmed by repeating the pupil-orofacial motion (PM) space projection at alternative micro-state counts K (**Fig. S5C**).

**Figure 4.**
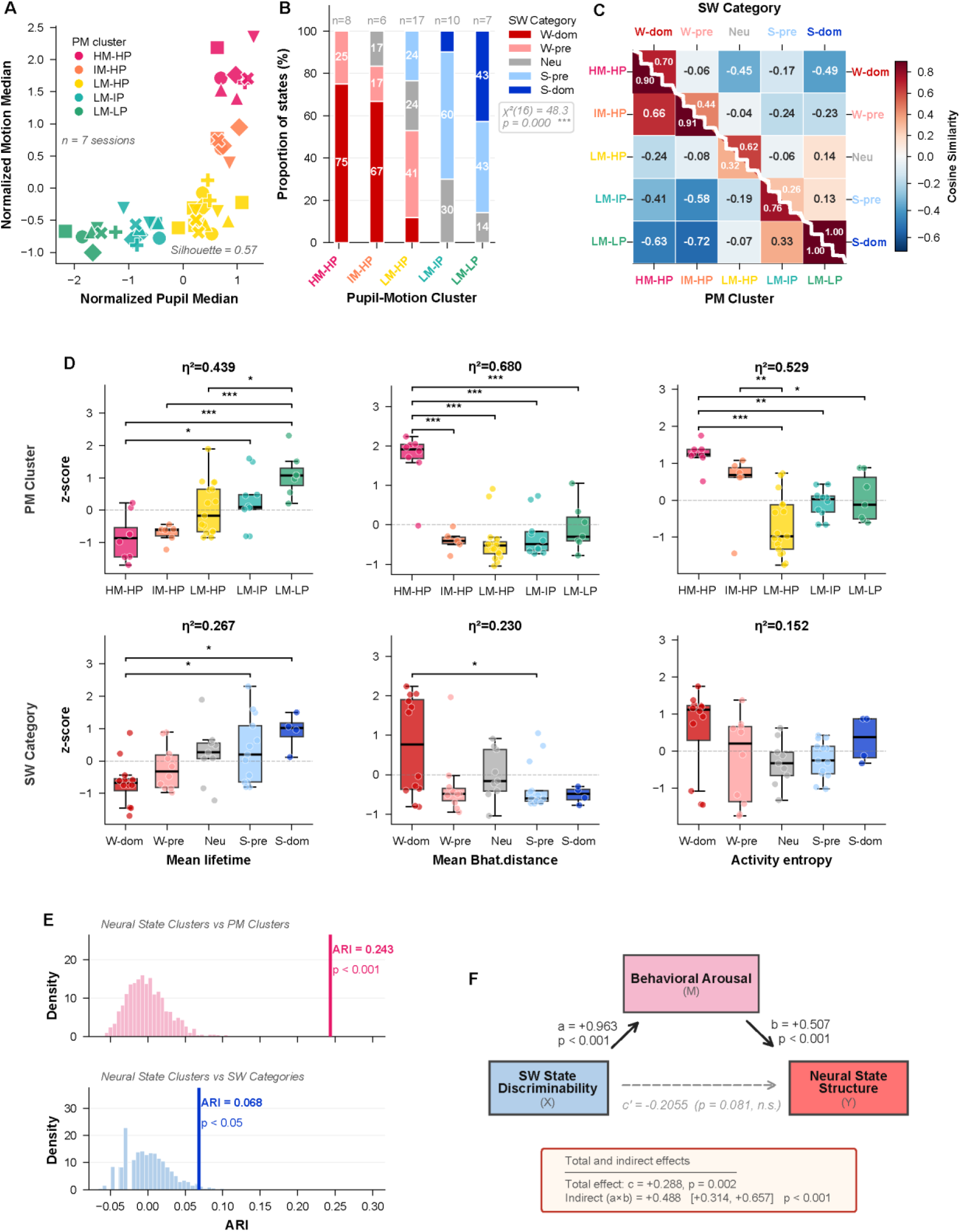
Neural micro-state structure is better captured by pupil-motion clustering than by sleep-wake categorization. **(A)** Pupil-Motion (PM) clustering of HMM micro-states across all sessions (n = 7). Each data point represents one micro-state, plotted by its median normalized pupil diameter (x-axis) and median normalized orofacial motion energy (y-axis). Five PM clusters were identified by k-means clustering: high-motion/high-pupil (HM-HP), intermediate-motion/high-pupil (IM-HP), low-motion/high-pupil (LM-HP), low-motion/intermediate-pupil (LM-IP), and low-motion/low-pupil (LM-LP). Animal identity is indicated by marker shape. Silhouette score = 0.57. **(B)** Proportion of HMM states assigned to each sleep-wake (SW) category within each pupil-motion (PM) cluster. SW categories are: Wake-dominant (W-dom), Wake-prefer (W-pre), Neutral (Neu), Sleep-prefer (S-pre), and Sleep-dominant (S-dom). Numbers above bars indicate total state count per PM cluster. Chi-square test of independence: χ²(16) = 48.3, p<0.001. **(C)** Composite cosine similarity heatmap of mean population activity vectors, averaged across sessions (n = 7). Lower triangle: group-averaged similarity between PM clusters; upper triangle: group-averaged similarity between SW categories. Diagonal values reflect within-group similarity. Color scale indicates cosine similarity. **(D)** Distributions of three neural micro-state features — mean lifetime, mean Bhattacharyya distance, and activity entropy — across PM clusters (top row) and SW categories (bottom row). Each data point represents one micro-state; box plots show median and interquartile range. Effect size (η²) and pairwise significance levels are indicated (*, p < 0.05; **, p < 0.01; ***, p < 0.001). **(E)** Adjusted Rand Index (ARI) quantifying the agreement between neural state clusters (k-means, neural feature space) and PM clusters (top) or SW category labels (bottom). Observed ARI values (vertical lines) are compared against a permutation null distribution (n = 1,000 permutations; shaded histograms). PM: ARI = 0.243, p < 0.001; SW: ARI = 0.068, p < 0.05. **(F)** Single-mediator model testing whether Behavioral Arousal (M) mediates the relationship between SW State Discriminability (X) and Neural State Structure (Y). Solid arrows indicate path a (X → M) and path b (M → Y | X); the dashed arrow indicates the conditional direct path c′ (X → Y | M). The inset reports the total effect, c, and the indirect effect, a×b. The indirect effect was evaluated by animal-level bootstrap resampling; its bootstrap distribution is shown in **Fig. S7F**.

To examine how PM clusters relate to conventional sleep-wake (SW) staging, we asked whether micro-states assigned to the same PM cluster shared similar SW fractional occupancy. Micro-states were assigned to SW categories by clustering them according to their sleep-wake discriminability (Cohen’s *d*, as in **Fig. 3D**) using k-means, yielding five ordered categories from Wake-dominant (W-dom) to Sleep-dominant (S-dom) (**Fig. S6A**). SW categories were not uniformly distributed across PM clusters (χ²(16) = 48.3, p < 0.001; **Fig. 4B**), indicating that SW category and PM cluster are related – micro-states within the same PM cluster are not uniformly distributed across SW categories – but also not redundant – PM and SW clustering are not fully overlapping. We therefore asked whether PM clusters or SW categories better captured the structure of population activity.

To this aim, we first computed the mean neuronal activity vector for each micro-state and quantified – separately for each recording session – the pairwise cosine similarity between micro-states. We then averaged these similarities within and across clusters to obtain cluster-by-cluster similarity matrices for PM clusters and SW categories respectively (**Fig. 4C**). PM clusters showed a wide, graded range of between-group similarity (–0.72 to 0.66), with behaviorally adjacent clusters showing moderately positive similarity (e.g., HM-HP vs. IM-HP: 0.66) and behaviorally distant clusters showing strongly negative similarity (e.g., HM-HP vs. LM-LP: –0.63). SW categories, in contrast, showed a narrower between-group range of similarity values (–0.49 to 0.14), with no strongly positive between-group similarity, indicating that SW categorization does not capture the full gradient of population activity structure. This pattern was robust to the number of HMM states used: repeating the cosine similarity analysis at alternative state counts K yielded a comparable between-group gradient for PM clusters but not for SW categories (**Fig. S5D**). Together, these results indicate that PM clustering captures a broader and more structured gradient of population activity than SW categorization. To further investigate this, we asked whether this difference extended to the temporal dynamic and neural activity structure of micro-states. We therefore compared three neural state features: mean lifetime, mean Bhattacharyya distance, and activity entropy (see **Methods**) across PM clusters and SW categories (**Fig. 4D**). All three neural state features differed across PM clusters, with large group effect sizes (η² = 0.439, 0.680, and 0.529 respectively), with HM-HP states showing the highest Bhattacharyya distance and activity entropy, that these high-arousal states were more separated from other micro-states in neural activity space and showed more heterogeneous population-activity profiles. In contrast, LM-LP states had the longest mean lifetime, indicating more prolonged persistence of this low-arousal neural configuration. The corresponding differences across SW categories were weaker (η² = 0.267, 0.230, and 0.152 respectively). SW categories did not significantly differ in activity entropy, and the significant differences in mean lifetime and Bhattacharyya distance were limited to a small number of comparisons between wake– and sleep-dominant categories. Together, these results indicate that pupil-motion clusters captured differences in the temporal and neural properties of micro-states more clearly than conventional sleep-wake categories.

To further characterize the neural feature space underlying micro-state identity, we applied k-means clustering independently of both PM and SW labels to the z-scored neural feature vectors, yielding five neural state clusters with distinct feature profiles (**Fig. S6E**). We quantified agreement between neural-cluster assignments and PM-cluster or SW-category labels using the Adjusted Rand Index (ARI), a chance-corrected measure of similarity between two partitions for which 1 indicates identical assignments and 0 indicates agreement expected by chance. Both label sets showed above-chance agreement with neural clustering relative to permutation null distribution (n = 1,000 shuffles; **Fig. 4E**), but agreement was higher for PM clusters (ARI = 0.243, p < 0.001) than for SW categories (ARI = 0.068, p < 0.05).

Nonetheless, the above-chance agreement of both PM clustering and SW categorization with neural state clustering raises the question of whether SW category and PM cluster provide distinct accounts of neural state structure, or whether the association between SW category and neural state structure reflects its correspondence with behavioral arousal. To address this question, we implemented a single-mediator linear mixed-effects mediation model (see **Methods**), in which the total effect of SW state discriminability on neural state structure is decomposed into a direct path and an indirect path via behavioral arousal (**Fig. 4F**). SW state discriminability, defined as the z-scored Cohen’s *d* for the difference in fractional occupancy between WAKE and NREM epochs, served as the independent variable (X). Behavioral arousal, summarized by the first principal component (PC1) of pupil diameter and orofacial motion energy, which explained 84% of their variance, served as the mediator (M; **Fig. S7A**). Neural state structure, summarized by the first principal component of mean lifetime, activity entropy, and mean Bhattacharyya distance, which explained 52% of the variance across these neural features, served as the outcome (Y; **Fig. S7B**). SW state discriminability strongly predicted behavioral arousal (path a = +0.963, p < 0.001, **Fig. S7C**), and behavioral arousal predicted neural state structure after controlling for SW state discriminability (path b = +0.507, p < 0.001, **Fig. S7D**). SW state discriminability was also positively associated with neural state structure overall (total effect c = +0.288, p = 0.002, **Fig. S7E**). The resulting indirect effect through behavioral arousal was significant (a×b = +0.488, 95% CI [+0.314, +0.657], p < 0.001, **Fig. S7F**). In contrast, the direct association between SW state discriminability and neural state structure after accounting for behavioral arousal was not significant (c′ = –0.206, p = 0.081), indicating that SW state influences neural state structure primarily through its association with behavioral arousal, with no significant independent contribution of SW state category.

### Distinct neuronal subpopulations show opposite tuning along the behavioral arousal axis

Having shown that the architecture of neural micro-states is explained by the PM-derived behavioral arousal axis, we next examined how the transitions between these micro-states are governed. We observed that the LM-HP cluster occupies a central bridging position in the state transition graph: while transitions between the higher-arousal clusters (HM-HP ↔ IM-HP) and between the lower-arousal clusters (LM-IP ↔ LM-LP) were frequent, direct transitions between the two ends of the arousal axis were rare, with LM-HP serving as the dominant intermediary between them (**Fig. 5A; Fig. S8**). At the population level, this arousal axis was mirrored by a systematic gradient in neural activity: higher-arousal states were characterized by few strongly activated neurons, while lower-arousal states engaged a distinct population of neurons, which was broader and less active, as reflected in the session-averaged activity fingerprint (**Fig. 5B**). Together, these observations suggest that the arousal axis is reflected in both the structure of state transitions and the organization of population activity. However, this does not reveal whether individual neurons consistently track position along the arousal axis across multiple micro-state transitions.

**Figure 5.**
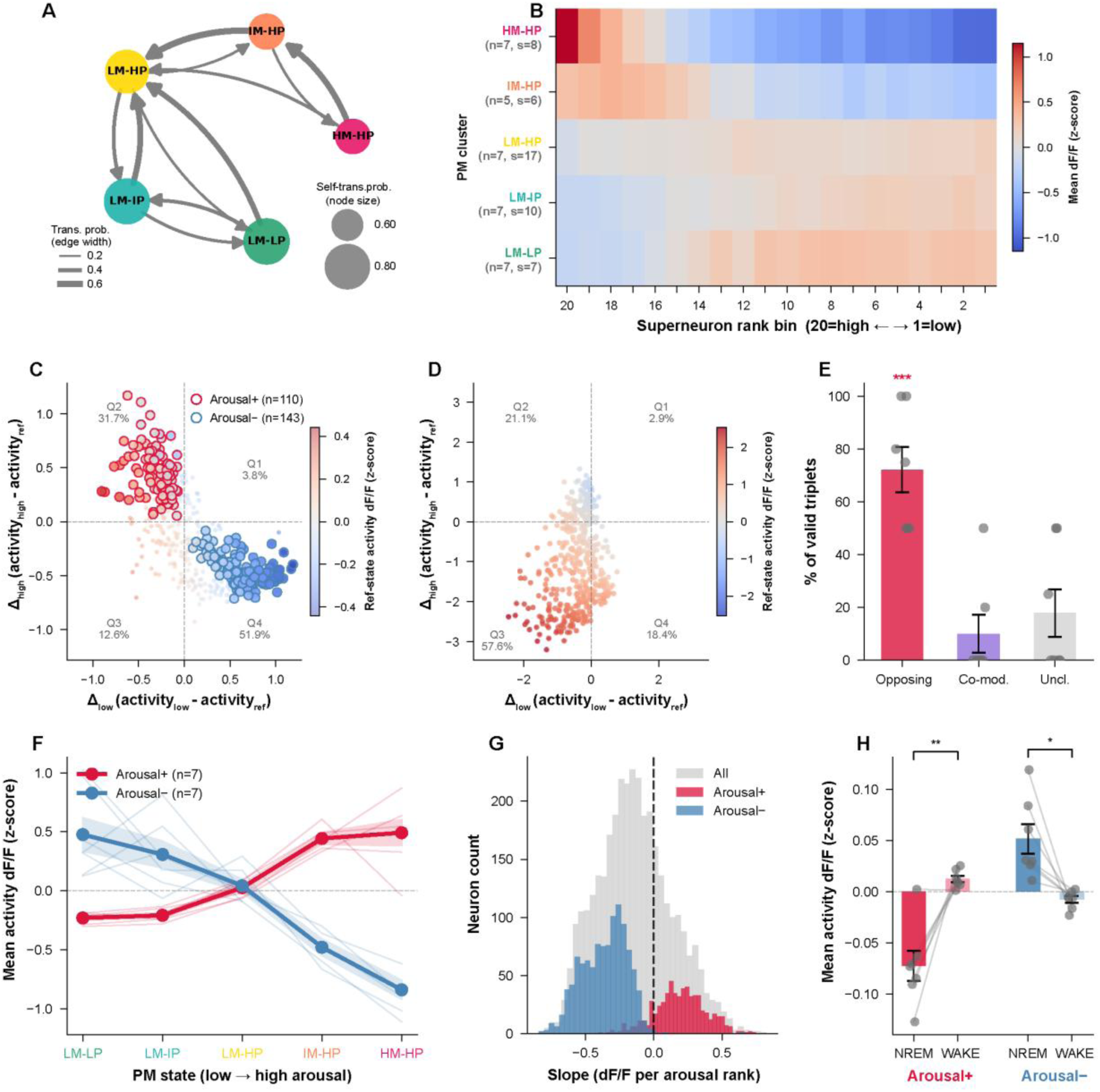
Distinct arousal-tuned neuronal ensembles track micro-state transitions in V1. **(A)** Directed transition graph of the five PM behavioral clusters, constructed from transition probabilities averaged across the seven imaging sessions. Node colors correspond to PM cluster identity; node size encodes self-transition probability (scale inset). Directed edges show inter-state transitions with probability > 0.2 (for visualization only); edge width is proportional to transition probability (scale inset). Full across-session transition matrices, including all transitions, are shown in **Fig. S8**. **(B)** Cross-session “population fingerprint” of neuronal activity across PM clusters. Each row represents one PM cluster (n = number of sessions; s = number of HMM states contributing to that cluster). Columns correspond to 20 percentile-rank bins of neuronal activity (bin 20 = most active, bin 1 = least active), where neurons are ranked within each session to equalize bin sizes across sessions. Color encodes mean dF/F (z-score) averaged across all contributing HMM states and sessions. **(C)** Scatter plot of one representative opposing triplet from an example session. Each point is one neuron. The x-axis shows the mean activity difference between the reference state and a lower-arousal state (Δ_low_ = activity_low_ − activity_ref_); the y-a xis shows the mean activity difference between the reference state and a higher-arousal state (Δ_high_ = activity_high_ − activity_ref_). Color indicates mean activity (dF/F, z-score) in the reference state. Quadrant labels report the percentage of neurons in each quadrant. Outlined circles mark neurons classified as Arousal+ ensembles (crimson, Q2; n = 110) or Arousal− ensembles (blue, Q4; n = 143), defined by intersection of quadrant assignments across all valid opposing triplets in this session. Dashed lines indicate zero on each axis. **(D)** Same as (C), for one representative co-modulated triplet from the same session. **(E)** Composition of triplet groups across sessions. Bars show the mean percentage of valid triplets classified as Opposing (crimson; 72.1 ± 8.6%), Co-modulated (purple; 10.0 ± 7.2%), or Unclassified (grey; 17.9 ± 9.0%) across 7 sessions (mean ± SEM); error bars denote SEM, and grey dots show individual sessions. Opposing fraction was tested against zero (one-sample t-test; *** p < 0.001). **(F)** Across-session mean activity trajectories of Arousal+ (crimson) and Arousal− (blue) ensembles across the five PM clusters ordered from low to high arousal (LM-LP → LM-IP → LM-HP → IM-HP → HM-HP). Thick lines show the cross-session mean (n = 7 sessions); shaded bands indicate ± SEM; thin lines show individual session trajectories. Dashed line indicates zero. X-axis tick colors correspond to PM cluster identity as in (A). **(G)** Distribution of linear-regression slopes for all recorded neurons (grey), Arousal+ ensembles (crimson), and Arousal− ensembles (blue), pooled across sessions. Slopes reflect the linear fit of each neuron’s mean activity (dF/F, z-score) across the five PM clusters ordered by arousal rank (1–5). The dashed vertical line indicates zero slope. Arousal+ and Arousal− ensembles showed large and opposing shifts in slope relative to the general population (Cohen’s *d* = 1.42 and −0.99, respectively; both p < 0.001, two-sample t-test). **(H)** Mean activity of Arousal+ (crimson) and Arousal− (blue) ensembles during NREM epochs and WAKE epochs across 7 sessions. Arousal+ ensembles: NREM = −0.072 ± 0.015, WAKE = 0.012 ± 0.003 (dF/F, z-score; paired t-test, p = 0.003). Arousal− ensembles: NREM = 0.052 ± 0.015, WAKE = −0.008 ± 0.003 (paired t-test, p = 0.015). Bars show mean ± SEM; grey dots represent individual sessions, with lines connecting paired values. Significance brackets: * p < 0.05, ** p < 0.01.

To address this, we defined state-transition triplets, each consisting of one higher-arousal state (HM-HP or IM-HP), LM-HP as the intermediate reference state, and one lower-arousal state (LM-IP or LM-LP). For each micro-state triplet, we calculated each neuron’s activity difference relative to the LM-HP reference state for the higher-and lower-arousal states. We then plotted the difference between the higher-arousal and reference states on the y-axis ( Δ_high_ = activity_high_ − activity_ref_), against the difference between the lower-arousal and reference states on the x-axis (Δ_low_ = activity_low_ − activity_ref_). This places each neuron in one of four quadrants (Q1-Q4, **Fig. 5C,D**).

Across micro-state triplets, two distinct scatter patterns of neuronal activity were observed: an anti-diagonal pattern in which neurons were distributed predominantly across Q2 and/or Q4, reflecting opposing activity changes in the two directions of arousal change (opposing triplets, **Fig. 5C**), and a main-diagonal pattern in which neurons mainly clustered in Q1 and/or Q3, reflecting activity changes of the same sign regardless of direction of arousal change (co-modulated triplets, **Fig. 5D**). Triplets were classified using the permutation-based quadrant criterion together with the across-neuron correlation between Δ_low_and Δ_high_ (**Methods; Fig. S9**). Across sessions, the majority of micro-state triplets were classified as opposing (72.1 ± 8.6%), while co-modulated (10.0 ± 7.2%) and unclassified (17.9 ± 9.0%) triplets were less common (**Fig. 5E**; **Fig. S9C–D**), indicating that bidirectional arousal tuning is the dominant organizational principle across the imaged population.

We next focused on opposing triplets to identify neurons with reproducible, oppositely directed activity changes across the arousal axis. Within each micro-state triplet, we retained Q2 and Q4 neurons only when their activity differences from LM-HP were significant for both the higher– and lower-arousal states. We then identified Arousal+ neurons as those retained in Q2 across all opposing triplets within a session, reflecting increased activity toward higher arousal and decreased activity toward lower arousal; Arousal− neurons were analogously identified from Q4 and showed the opposite profile. This yielded two session-level ensembles: Arousal+ ensembles (16.7 ± 4.1% of recorded neurons) and Arousal− ensembles (Q4; 32.2 ± 5.6% of recorded neurons; **Fig. S9E**).

We then characterized the activity profiles of Arousal+ and Arousal-ensembles across the PM-derived behavioral arousal gradient. When PM clusters were ordered from low to high arousal (LM–LP → HM–HP), Arousal+ ensembles showed progressively increasing activity, whereas Arousal− ensembles exhibited the opposite pattern, with activity decreasing across the arousal gradient (**Fig. 5F**). Consistent with this tuning, the distribution of arousal-related slopes in ΔF/F values was significantly biased toward positive values for Arousal+ ensembles and toward negative values for Arousal− ensembles, relative to the general population of neurons (**Fig. 5G**). To confirm that this tuning extended beyond discrete PM states, we examined ensemble activity as a continuous function of pupil diameter and orofacial motion: Arousal+ ensembles showed increasing activity with larger pupils and stronger orofacial motion, whereas Arousal− ensembles showed the opposite (**Fig. S10A-B**). At the macro-state level, Arousal+ ensembles were more active during wakefulness than NREM sleep, whereas Arousal− ensembles showed the opposite pattern (**Fig. 5H**), in line with the correlated nature of macro– and micro-states along a pupil–orofacial arousal axis. Finally, despite their opposing functional profiles, Arousal+ and Arousal− ensembles were spatially intermingled, with centroids in close proximity and only modest differences in mean pairwise distance between across versus within classes (**Fig. S10C-D**), indicating no clear large-scale spatial segregation of the two neuronal subpopulations.

Together, these results demonstrate that V1 contains two distinct neuronal subpopulations with opposing and consistent tuning along the arousal axis: Arousal+ ensembles that increase activity with rising arousal and are preferentially active during wakefulness, and Arousal− ensembles that show the opposite profile and are more active during NREM sleep. This opponent organization suggests that arousal shapes V1 activity through the differential and coordinated recruitment of distinct neuronal subpopulations, rather than a uniform gain modulation of the imaged population.

## Discussion

Using large field-of-view two-photon calcium imaging in head-fixed mice, we characterized the organization of V1 population dynamics across the sleep-wake cycle. We show that V1 population activity is not fully characterized in terms of discrete sleep-wake categories. Instead, activity consistently entered temporally stable micro-states characterized by distinct population-activity patterns and organized transitions between micro-states. Although individual micro-states varied in their relative frequency during NREM and wakefulness, none was exclusive to either macro-state, and their neural activity properties were more closely aligned with a continuous behavioral arousal axis derived from pupil diameter and orofacial motion than with the conventional NREM-WAKE distinction. At the circuit level, this organization was reflected in two oppositely tuned neuronal ensembles: Arousal+ and Arousal− neurons that showed, respectively an increase and a decrease in activity across the behavioral arousal gradient. Together, these findings suggest that behavioral arousal provides a continuous organizing dimension for cortical population dynamics that is not fully captured by conventional sleep-wake categorization.

While circadian and homeostatic factors are major determinants of large-scale oscillatory brain dynamics^12,31,32^, our findings thus suggest that behavioral arousal provides an additional, and possibly equally important, organizing dimension at the level of neuronal population activity. Previous work showed that pupil-linked fluctuations in arousal track rapid changes in cortical state and shape both spontaneous and sensory-evoked activity in awake V1^17,18,33^. More broadly, spontaneous behavior provides important structure to brain-wide population activity^34^, and arousal has more recently been proposed as a low-dimensional determinant of widespread cortical dynamics^35^. These studies, however, have largely focused on wakefulness. Our results extend the question of arousal-linked population organization into NREM sleep, showing behavioral arousal tracks the organization of V1 micro-states across conventional sleep-wake categories.

Our identification of recurring V1 micro-states suggests that cortical population activity is organized at a finer temporal and dynamical scale than conventional sleep-wake labels capture. This upends the classical subdivision of brain activity patterns into a limited set of uniform, non-overlapping brain states^11^, but is nonetheless consistent with evidence that cortical activity occupies a constrained repertoire of recurring population-levels configurations^36^ and that cortical networks can display local, short-lasting patterns of activity that transiently diverge across brain regions^14,16,23,37,38^. Importantly, our findings significantly extend this perspective by showing that V1 micro-states are also embedded within a structured transition topology. Population activity preferentially progressed between micro-states occupying nearby positions along the behavioral arousal axis, rather than directly switching between states associated with low– and high-arousal extremes – and also with NREM sleep and wakefulness, respectively. This pattern parallels HMM-based studies of human whole-brain dynamics, in which recurrent brain states also exhibit structured and non-random transition patterns^39,40^. Altered frequency and transition dynamics of recurrent brain states have also been reported in narcolepsy and associated with clinical symptoms, raising the possibility that micro-state-dynamic measures could potentially provide quantitative markers of unbalanced state regulation^41^. The persistence of this organization across both wakefulness and sleep further indicates that micro-state transition pathways reflect intrinsic properties of V1 population dynamics.

At the circuit level, the opposite tuning of Arousal+ and Arousal− neurons may provide a cellular substrate for the behavioral arousal axis in V1. This parallels recent evidence for two analogous, oppositely modulated neuronal populations distributed across multiple brain regions and associated with widespread hemodynamic fluctuations^42^. The cross-regional occurrence of analogous opponent neuronal populations, together with evidence that arousal-related dynamics shape large-scale brain activity, therefore suggests that arousal-linked population organization may extend beyond V1^35,42^. The spatial intermingling of Arousal+ and Arousal-neurons further suggests that this architecture can be implemented among functionally distinct neurons without requiring anatomically segregated modules. However, the functional role of these opposing populations remains unresolved. Their opposite tuning may primarily reflect common modulation by an arousal-related signal, or it may contribute to the stabilization, reconfiguration, or transition of local population micro-states. Causal studies in sleep-regulatory circuits show that neuronal populations recruited during prolonged wakefulness can actively promote subsequent NREM-sleep transitions^43^. Arousal+ and Arousal− neurons in V1 might provide a substrate for this phenomenon, a hypothesis that will have to be tested via ensemble-specific perturbations combined with population-level recording. In V1, furthermore, arousal dynamically changes visual response gain and population coding, suggesting that the population micro-state occupied at a given moment may influence how sensory information is processed^18^. More broadly, fine-scale micro-state dynamics may similarly shape offline processing. Recent studies have shown that sleep microstructure can organize distinct memory-replay processes^44^ and targeted memory reactivation at specific phases of ongoing neural oscillations – possibly reflecting the presence of micro-states – during sleep can influence memory consolidation^45–47^. Although these studies do not establish a role for the V1 micro-states identified here, they motivate the testable hypothesis that cortical micro-states and their transitions are functional population configurations that gate or shape sensory processing during wakefulness and offline memory processing during sleep.

Several limitations need to be acknowledged. First, recordings were obtained in head-fixed mice, a preparation that constrains natural behavior and can alter aspects of sleep architecture^48^. In addition, REM sleep was infrequent in our recordings. Neuronal assembly composition has been shown to reorganize across NREM–REM sequences^49^, raising the possibility that REM sleep engages a distinct mode of population organization not captured by the framework described here. Second, although large-scale two-photon imaging enabled tracking of individual layer II/III intratelencephalic pyramidal neurons in V1 – providing evidence that the micro-state architecture we observed is functionally rather than genetically determined – its temporal resolution limits the characterization of rapid activity patterns and transition dynamics. Moreover, it remains unclear whether the observed micro-states and Arousal+/Arousal− ensembles generalize to other neuronal classes, cortical layers, or brain regions. Finally, our findings are correlational. The pupil– and orofacial-related arousal measure statistically accounted for variation in micro-state structure, but neither this association nor the mediation analysis establishes that behavioral arousal causally drives micro-state transitions. Resolving the underlying temporal and causal mechanisms will require simultaneous electrophysiological recordings and targeted perturbation of arousal-related systems or activity-defined ensembles.

Overall, our study establishes a framework in which the nature of V1 population activity is determined by and organized through recurring micro-states and structured transitions along a continuous behavioral arousal dimension, rather than by discrete sleep-wake categories. This challenges the view of brain states as discrete configurations that determines cortical activity at a global – and sometimes local – level, and instead suggests that what we call brain state may emerge from the continuous, behavioral arousal-linked organization of local population dynamics across the sleep-wake cycle.

## Methods

### Animals

All animal experiments were approved by the Dutch Commission for Animal Experiments (CCD application number: AVD11100202215932) and by the Animal Welfare Body of the University of Amsterdam. Mice were housed under a 12-hour light/dark cycle with ad libitum access to food and water. Home cages were equipped with bedding and a running wheel to promote locomotor activity during the dark phase. All recordings were performed during the light phase, with each session lasting no more than 3 hours starting from lights onset.

Mice were double-transgenic offspring generated by crossing the Rasgrf2-dCre driver line (JAX #022864) with the Ai148D reporter line (JAX #030328), which together drive GCaMP6f expression selectively in a subset of layer II/III cortical excitatory neurons corresponding to the intratelencephalic (IT) projection class^50,51^, via a trimethoprim (TMP)-inducible destabilized Cre (dCre) recombinase^52^ and the TIGRE2.0 reporter system^53^. This transgenic cross has been widely used for in vivo two-photon and functional cortical calcium imaging^54,55^. In the absence of TMP, dCre undergoes proteasomal degradation, preventing recombination. To induce GCaMP6f expression, mice received intraperitoneal TMP injections (0.25 mg/g) on three consecutive days, at least one week before surgery. Experiments were conducted on adult mice (2–6 months old).

For two-photon calcium imaging (Dataset A), data were collected from n = 7 mice (5 males, 2 females). A subset of these animals (n = 2 males) subsequently underwent Neuropixels recordings (Dataset B), conducted at least one month after the final two-photon imaging session to allow full recovery from any possible cumulative effects of chronic imaging.

### Surgical procedures

Mice were anesthetized with isoflurane (5% induction, 1.5–2% maintenance) and placed in a stereotaxic frame with body temperature monitored. To minimize brain oedema, dexamethasone (8 mg/kg, intramuscular) was administered prior to incision. Following scalp disinfection and incision, the skull surface was cleaned and dried. A custom titanium head bar was affixed to the skull using cyanoacrylate adhesive and dental cement to enable subsequent head fixation.

A large craniotomy encompassing the entire left hemisphere was performed, under periodic saline irrigation to prevent tissue dehydration. The skull was carefully removed, and the exposed brain was flushed with cold, filtered saline. A custom polydimethylsiloxane (PDMS) cranial window was placed over the craniotomy and sealed with surgical adhesive, following established procedures^56,57^. Dental cement was applied around the head bar and window perimeter to stabilize the implant.

Following surgery, mice received pain-relief-enriched water (Carprofen and Baytril in sucrose solution) and were housed in cages with soft bedding. Window optical quality was assessed daily over a two-week post-surgical recovery period; mice with suboptimal window quality were excluded before proceeding to habituation.

### Habituation to the experimental setup

Following a minimum of two weeks of post-surgical recovery, mice were gradually accustomed to head fixation in the experimental setup to facilitate the natural emergence of sleep under the microscope. Habituation sessions were conducted under the same environmental conditions as actual recordings — in darkness and at ambient noise levels, and beginning 1 hour after light onset to allow animals to complete their initial sleep period before handling — with the exception that the imaging laser was not active. Session duration was progressively increased over seven days, from 15 minutes on day 1 to 3 hours by day 7. Mice were subsequently head-fixed for 3 hours per day for three additional days immediately prior to the start of recordings.

### Retinotopic mapping

Before two-photon imaging, widefield calcium imaging was used to identify the location of primary visual cortex (V1) within the cranial window. The cortical surface was illuminated with a 470 nm LED (Thorlabs M470L5, driver: Thorlabs T-Cube) and imaged through a 1.25× objective (Leica HC PL FLUOTAR 1.25×/0.04 T) with a bandpass filter cube (Leica I3 DM 513828) and a CMOS camera (Basler ace 2 a2A1920-160um).

Moving checkerboard-bar stimuli were presented to the contralateral right eye in four directions, with 20 repetitions per direction^54^. Retinotopic maps were generated using custom Python code adapted from the phase-encoded mapping approach described by Zhuang et al.^58^ and implemented as described by Bassler et al.^54^. Briefly, responses were averaged across trials for each movement direction, and Fourier analysis at the bar-sweep frequency was used to generate phase maps. Phase maps from opposing directions were combined to obtain azimuth and altitude maps, which were smoothed with a two-dimensional Gaussian filter. Gradients of the azimuth and altitude maps were then combined to generate a visual field-sign map. V1 and neighboring visual areas were identified as contiguous regions with shared field-sign values. The resulting map was used to position the two-photon imaging field of view within V1.

### Two-photon calcium imaging (Dataset A)

Head-fixed mice were imaged under a large field-of-view two-photon microscope (Bergamo II, Thorlabs) equipped with a 16× 0.8 NA objective (Nikon) and a digital zoom of 2×, yielding a field of view of 973.54 × 973.54 µm. Imaging was performed using an 80 MHz ultrafast pulsed laser at 920 nm (Spark Lasers) with 38.3 mW power at the sample, and fluorescence was detected using PMT detectors. Calcium signals were recorded at a single focal plane approximately 150–300 µm below the cortical surface. Frames were acquired at ∼7.7 Hz (1,024 × 1,024 pixels; 60,000 frames per session, corresponding to ∼2 hours and 10 minutes) using ThorImageLS 4.3 software. Recordings were conducted in darkness with no visual stimulation to allow spontaneous alternation between wakefulness and sleep. We planned up to two recording sessions per mouse. Further longitudinal sampling was not pursued because each session required approximately one week of habituation to the head-fixed recording setup, and the feasible recording period was limited by post-surgical window quality and animal age. In total, we acquired 10 sessions from 7 mice. Not all mice completed a second session because of declining cranial-window quality or other technical problems. Three acquired sessions were excluded from further analysis: two due to an insufficient number of NREM epochs for reliable estimation of session-level micro-state features, and one due to visible ocular discharge during a second recording session in the same animal, which compromised pupil tracking. The final dataset comprised n = 7 sessions from 7 mice.

### Neuropixels recording (Dataset B)

Extracellular electrophysiological recordings were performed in a subset of animals (n = 2 mice, 3 sessions per animal) approximately one month after the final two-photon imaging session. A small circular incision (1.5 mm diameter) was made in the PDMS cranial window directly above V1, as identified by prior retinotopic mapping, to allow acute probe insertion^57^. A Neuropixels 1.0 probe (IMEC) was inserted vertically up to a depth of ∼2000 µm from the brain surface using a micromanipulator. Between sessions, the incision was resealed with PDMS and silicone gel to protect the opening on the cranial window for repeated access.

LFP signals were acquired at 2500 Hz via the dedicated LFP stream in Open Ephys (v0.6.6). Raw traces were phase-shift corrected to account for inter-channel hardware filter delays inherent to Neuropixels 1.0, using the *phase_shift* function implemented in SpikeInterface^59^. For subsequent analysis, LFP signals from channels C130–C180 were used, corresponding to approximately 200–700 µm below the cortical surface based on the known inter-channel spacing of 20 µm (IMEC Neuropixels 1.0 specifications). Further processing is described in the *Brain state scoring section from LFP* section. Recording sessions were conducted under the same environmental conditions as two-photon imaging sessions — in darkness and without visual stimulation — to allow spontaneous alternation between wakefulness and sleep.

### Acquisition of pupil area and orofacial motion

In both two-photon imaging and Neuropixels recording sessions, pupil area and orofacial motion were recorded simultaneously with neural data using a digital infrared camera (Basler GigE) at ∼30 frames per second, capturing 1282 × 1026-pixel images. During two-photon imaging sessions, the two-photon laser provided sufficient infrared illumination for pupil imaging^27^. During Neuropixels recording sessions, a dedicated IR-LED (860 nm) was positioned above the cranial window, adjacent to the probe, to illuminate the eye. Synchronization with neural recordings was achieved via a digital TTL signal, with timestamps used for post hoc alignment of pupil and orofacial motion data to the neural recording timeline.

### Brain state scoring from LFP

Brain states were scored manually from LFP recordings (Dataset B). Raw LFP signals were downsampled to 1000 Hz and referenced against the common average of all recorded channels (C1–C191). The average LFP trace across channels C130–C180 was segmented into non-overlapping 10-second epochs, each visually inspected and labelled as NREM sleep or Wakefulness (hereafter WAKE). NREM epochs were identified by the presence of clear slow-wave oscillations (0.5 **–** 4 Hz); WAKE epochs were characterized by a comparatively flat, low-amplitude LFP ^2,9^. Orofacial motion was used as a reference to aid state disambiguation. Epochs that did not meet the criteria for either NREM or WAKE — including epochs with low-amplitude LFP but absent slow-wave oscillations, and low orofacial motion, which may include REM sleep — were labelled as Unclassified and excluded from all subsequent analyses. As electromyography (EMG) was not recorded, unambiguous identification of REM sleep was not possible and it was therefore not treated as a separate category. All epochs were scored independently by at least two raters; discrepancies were resolved by consensus discussion.

### Pupil & orofacial motion feature extraction

Raw video frames were inverted prior to analysis using FFmpeg, as the infrared illumination renders the pupil bright against a dark background. Pupil diameter and orofacial motion energy were then extracted from the inverted video using FaceMap^24^. Raw pupil area was converted to pupil diameter by taking the square root of the area per frame, which was then normalized per session by dividing it by the mean of the top 100 frames^27^. To extract features for brain state classification, normalized pupil diameter traces were segmented into 60-second epochs with a 10-second sliding step. This epoch length was chosen to capture the slow temporal dynamics of pupil fluctuations associated with NREM, while remaining short enough to resolve state transitions^26,60^. For each epoch, four features were computed:

1. **Median pupil diameter**: the median normalized pupil diameter within the epoch
2. **Distribution width**: the difference between the 97.5th and 2.5th percentiles of the pupil diameter within the window ^27^
3. **Mean slope**: the mean of the frame-to-frame differences of the pupil diameter trace within the epoch, capturing the average rate of change
4. **Median lag-1**: the median pupil diameter of the immediately preceding 60-second epoch, providing temporal context across consecutive windows

These four features were used as inputs to the logistic regression classifier (see *Pupil-based brain state classification*). Occasional blinks, visible as brief transient drops in pupil diameter (**Fig. 2E**), were not explicitly removed prior to feature extraction; given their transient nature relative to the 60-second epoch length, blinks are unlikely to substantially affect epoch-level median pupil diameter or median lag-1 features, and their potential contribution to distribution width and mean slope was considered negligible given the rarity of blinks within any single epoch.

Orofacial motion energy was normalized per session by dividing by a baseline defined as the median of the lowest 5% of motion energy values^24^, and was used as a qualitative reference during manual LFP scoring (see *Brain state scoring from LFP*) but was not included as a feature for the classifier.

### Pupil-based brain state classification

To enable brain state classification in Dataset A (where no LFP was recorded), a logistic regression classifier (scikit-learn v1.4.0) was trained on Dataset B using LFP-derived NREM/WAKE labels as ground truth and pupil-based features as predictors. Prior to model fitting, all four pupil-derived features were standardized to zero mean and unit variance using a StandardScaler.

To align LFP-derived state labels with the 60-second pupil feature epochs, a sliding window approach was applied: a 60-second epoch was assigned a given state label only if at least 5 out of 6 consecutive 10-second LFP-scored epochs carried that label, ensuring temporal consistency. Epochs not meeting this criterion were labelled as Unclassified and excluded from classifier training and evaluation.

Model performance was evaluated using leave-one-session-out cross-validation, in which each session in Dataset B served as the test set once, while the remaining sessions were used for training; feature scaling was fitted on the training fold only to prevent data leakage. Classifier performance was assessed using the area under the receiver operating characteristic curve (AUC).

To ensure high-confidence state assignments, a posterior probability threshold was selected as the lowest threshold at which NREM precision reached ≥ 90%, as determined programmatically from the precision–recall curve (**Fig. 2C**), corresponding to P(NREM) ≥ 0.72. Epochs were labelled NREM if P(NREM) ≥ 0.72, WAKE if P(NREM) ≤ 0.28 (the symmetric counterpart, 1 − 0.72); epochs with intermediate probabilities (0.28 < P(NREM) < 0.72) were considered low-confidence and excluded from downstream analysis.

Following cross-validation, a final model was trained on all sessions of Dataset B to maximize the amount of training data for inference. This final model was then applied to each session of Dataset A independently, using the same feature extraction pipeline and the scaler fitted on the full Dataset B training set.

### Two-photon data preprocessing & Rastermap visualization

Two-photon calcium imaging data were processed using Suite2p^61^, which performed motion correction, region of interest (ROI) detection, and signal extraction. Putative neurons were identified using Suite2p’s built-in classifier and further refined by manual inspection of ROI masks and activity traces; only ROIs classified as cells were retained for downstream analysis.

The corrected fluorescence signal was computed as:

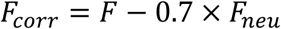

where F is the raw fluorescence, F_neu_ is the neuropil fluorescence, and 0.7 is the default neuropil contamination coefficient^61^.

The normalized fluorescence change (ΔF/F) was then calculated as:

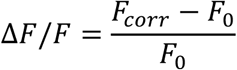

where *F_0_* is the baseline fluorescence estimated using a 60-second sliding window 10th-percentile filter applied to (*F_corr_*) with mirror-mode padding at the edges.

To visualize population-level activity patterns across brain states, the full-session ΔF/F matrix (neurons x time frames) was given as input to Rastermap^25^, run independently for each session. Rastermap embeds neurons into a 1D ordering based on the correlation structure of their activity, grouping neurons with similar temporal dynamics. For visualization, neurons were averaged into superneurons^25^ — bins of 5 neurons with similar temporal dynamics — to provide a smoothed overview of population activity. Superneurons were used for visualization purposes only and were not used in any downstream analyses; all subsequent analyses were performed on individual neuron ΔF/F traces.

### HMM micro-states analysis

To identify discrete population activity micro-states, a Gaussian Hidden Markov Model (HMM) with diagonal covariance — chosen to reduce the number of free parameters given the high-dimensional input — was fitted to the full-session ΔF/F matrix (T timepoints x N neurons) independently for each of the 7 sessions using the *hmmlearn* Python package. For each session, HMMs were fitted across K = 2–20 states. Each K was fitted with 5 independent random initializations to mitigate convergence to local optima, with a maximum of 200 Expectation-Maximization (EM) iterations and a convergence tolerance of 1×10⁻⁴ on the relative change in log-likelihood. For each value of K, solution stability was first quantified as the mean pairwise normalized mutual information (NMI) across all 5 initializations; the run with the highest log-likelihood (LL) was retained^62^.

### Per-session candidate pool and upper bound

For each session, a candidate pool of valid K values was constructed by applying two biological constraints to eliminate degenerate solutions: (1) minimum fractional occupancy ≥ 2% per state ^63^, ensuring each state accounted for a meaningful proportion of recording time; (2) minimum mean dwell time ≥ 2 frames (∼260 ms) per state^64,65^, ensuring states reflected temporally sustained activity patterns rather than transient fluctuations (**Fig. S3D–E**). K < 3 was excluded on modelling grounds as the trivial binary-switch solution. An upper bound K_up_ was then determined from the variance elbow — the point of maximum curvature in the average within-state variance curve, identified using the kneedle algorithm^66^ (**Fig. S3B**) — marking where additional states ceased to meaningfully reduce within-state heterogeneity. The log-likelihood elbow — the first K at which the marginal LL gain dropped below 20% of the K = 2→3 gain — was computed as a convergent indicator of diminishing model fit returns^62^ (**Fig. S3A**). Where the NMI stability ceiling reflected a substantial, persistent drop — absolute NMI drop > 0.02 and relative drop > 2% relative to the preceding K, not recovered to ≥ 90% of the prior value at K+1 — it was used to tighten K_up_ further^67,68^ (**Fig. S3C**). The candidate pool comprised all valid K values at or below K_up_ (**Fig. S3F**).

### Cross-session selection of the number of micro-states

K_up_ was not treated as a hard exclusion threshold but as an estimate of the upper limit of single-session informativeness. The final K per session was determined by cross-session optimization, which constitutes a stronger validation criterion: a K value yielding reproducible state structure across independently recorded sessions provides evidence that the additional states reflect genuine population dynamics rather than session-specific noise, even if that K exceeds what single-session criteria alone would support.

The cross-session search range was anchored to the distribution of per-session K_up_ values (K ∈ {5, 6, 7, 8}; **Fig. S3F**). An exhaustive search was performed over all K combinations within this range. Cross-session consistency was quantified as the mean pairwise Pearson correlation of Hungarian-matched state feature profiles across all session pairs^29^. Each state was characterized by four features: fractional occupancy (proportion of time spent in each state), self-transition probability (probability of remaining in the same state), activity entropy (Shannon entropy of the mean state emission vector), and population synchrony (mean pairwise Pearson correlation between neurons across all frames assigned to that state) — each z-scored independently across states within each session prior to matching. The K combination maximizing mean pairwise cross-session correlation was selected as the final session-specific state-number assignment (mean pairwise correlation = 0.80; **Fig. S3G–I**).

### Micro-state characterization

Following final K selection, each state was characterized by its behavioral and neural signatures. States were sorted by ascending mean population activity, and this ordering was preserved throughout all subsequent analyses. Pupil diameter and orofacial motion energy were aligned to the HMM frame timestamps by linear interpolation, as behavioral signals were acquired at a different sampling rate than the imaging data. State occupancy was computed as the percentage of frames assigned to each state within each high-confidence behavioral epoch (WAKE or NREM), as classified by the logistic regression classifier. State transition probabilities were obtained from the HMM transition matrix, estimated during model fitting from the inferred frame-to-frame state transitions.

### Pupil-motion (PM) clustering

For each HMM micro-state in each recording session, the median normalized pupil diameter and median normalized orofacial motion were computed across all frames assigned to that state. To remove between-session differences in baseline pupil size and movement level, these per-state medians were z-scored within each session (subtracting the session mean and dividing by the session standard deviation across all states in that session) before pooling across sessions.

Micro-states were clustered in the resulting two-dimensional PM feature space using k-means (*scikit-learn KMeans*; 50 random initializations, Euclidean distance). The number of clusters was set to K = 5, selected on the basis of the silhouette score evaluated across candidate solutions K = 2–6, which reached a maximum at K = 5 **(Fig. S5 A-B)**. Clusters were semantically labelled according to their centroid positions: the three clusters with the highest pupil centroid values were ranked by motion level and labelled HM-HP (high-motion/high-pupil), IM-HP (intermediate-motion/high-pupil), and LM-HP (low-motion/high-pupil); the two remaining clusters were ranked by pupil level and labelled LM-IP (low-motion/intermediate-pupil) and LM-LP (low-motion/low-pupil).

### Sleep and wakefulness (SW) category assignment

Sleep/wake dominance categories were assigned by computing, for each micro-state, Cohen’s *d* of fractional occupancy between WAKE and NREM epochs (positive values indicating WAKE preference, negative values indicating NREM preference). These per-state Cohen’s d values were then clustered using k-means (K = 5, matching the PM solution), and the resulting clusters were ordered from most Wake-dominant to most Sleep-dominant, yielding five categories: Wake-dominant (W-dom), Wake-prefer (W-pre), Neutral (Neu), Sleep-prefer (S-pre), and Sleep-dominant (S-dom).

### Population activity similarity

Pairwise cosine similarity was computed between the mean ΔF/F activity vectors of all micro-state pairs within each recording session. The mean activity vector of each HMM micro-state was defined as the state emission mean across all recorded ROIs. Cosine similarity between states i and j was computed as:

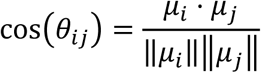

where μ_i_ denotes the mean ΔF/F activity vector of state i. Each micro-state was assigned a PM cluster label and a SW category label based on the behavioral data recorded in the same session. Pairwise cosine similarities were averaged across all state pairs sharing the same group combination and pooled across sessions, yielding session-averaged 5×5 group-by-group similarity matrices for PM clusters and SW categories respectively. These two matrices are displayed as a composite split-diagonal heatmap (**Fig. 4C**), with the lower triangle showing PM cluster similarity and the upper triangle showing SW category similarity.

### Neural feature-space clustering

#### Feature extraction and selection

Ten candidate features were initially extracted for each micro-state across three categories — temporal dynamics, state separation, and population-activity structure — to comprehensively characterize micro-state identity from complementary perspectives: temporal dynamics capture how long and how stereotypically a state recurs; state separation quantifies how distinctly a state occupies the neural activity space relative to other states; and population-activity structure describes the internal structure of population activity within a state^69,70^.

**Temporal dynamics** — reflecting how states evolve and persist over time: fractional occupancy, mean lifetime, self-transition probability, transition entropy^70–72^.

**State separation** — reflecting how distinctly a state occupies neural activity space: mean Bhattacharyya distance, minimum Bhattacharyya distance, Log-determinant of covariance^73^.

**Population-activity structure** — reflecting the pattern of population activity within a state: activity entropy, mean cosine similarity, population synchrony^74^.

To identify a compact, non-redundant feature set, pairwise Spearman correlations were computed between all ten features across all states and sessions. Features were then grouped by similarity (defined as 1 − |Spearman ρ|) using Ward linkage hierarchical clustering^75^, such that features carrying redundant information were placed in the same cluster^76^. Visual inspection of the resulting dendrogram identified three natural feature clusters (**Figure S6B**). To select the most informative representative from each feature cluster, we used the one-way ANOVA F-statistic as an objective criterion, as it directly quantifies how well each feature discriminates between neural states (between-state variance relative to within-state variance). Concretely, a preliminary K-means solution (K = 5) was fitted to all ten candidate features to provide cluster labels (similar to *Neural feature-space clustering*), and the feature with the highest F-statistic within each cluster was retained as its representative (**Figure S6C**).

This yielded three retained features:

1. **Mean lifetime:** the mean duration of consecutive frames assigned to a state, reflecting state temporal stability.
2. **Mean Bhattacharyya distance**: the mean pairwise Bhattacharyya distance between a state and all other states, reflecting state isolation in neural activity space. For two Gaussian states N(*μ_i_*, Σ*_i_*) and N(*μ_j_*, Σ*_j_*), the Bhattacharyya distance is:

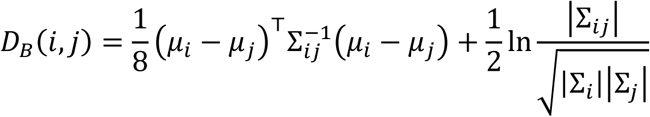 Where 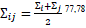
3. **Activity entropy:** the Shannon entropy of the absolute mean ΔF/F activity profile across neurons, computed as −Σ p·log(p) where p is the normalized absolute weight of each neuron. High values reflect diffuse activity distributed across many neurons; low values reflect activity concentrated in a small subset.

All three features were z-scored within each session prior to clustering and comparison.

#### Neural feature-space clustering

Micro-states were clustered in the three-dimensional neural feature space using K-means (K = 5, matching the number of PM clusters; 10 random initializations, random seed = 42). This yielded a set of neural feature-space cluster labels used as the reference partition for ARI computation.

### Neural feature characterization

To characterize how neural micro-state features varied across grouping schemes, group differences in each retained feature across PM clusters and SW categories were assessed using a one-way ANOVA (*scipy.stats.f_oneway*). Effect size was quantified as eta-squared (η² = SS_between_ / SS_total_, where SS_between_ is the sum of squares associated with differences among groups, and SS_total_ is the total sum of squares across all HMM micro-states included in the analysis. Thus, η² indicates the proportion of variation in the neural feature associated with group membership. Where significant (p < 0.05), post-hoc pairwise comparisons were performed using Tukey’s HSD test (*statsmodels pairwise_tukeyhsd*, α = 0.05).

### Adjusted Rand Index (ARI) and permutation testing

To quantify the alignment between neural feature-space structure and each behavioral cluster scheme, the ARI was computed between the neural feature-space cluster labels and the PM cluster labels, and separately between the neural feature-space cluster labels and the SW category labels. The ARI corrects for chance-level agreement and ranges from 0 (no agreement beyond chance) to 1 (perfect agreement)^79^.

Statistical significance was assessed by a permutation test (1,000 permutations, independent random seeds per arm), in which the external labels (PM or SW) were randomly permuted while preserving cluster sizes, generating a null ARI distribution. A permutation p-value was computed as the proportion of null ARIs exceeding the observed value; significance was inferred at p < 0.05.

### Mediation Analysis

To test whether arousal mediates the relationship between sleep/wake (SW) state discriminability and neural state structure, we implemented a single-mediator mediation model with the following variable mapping:

- **X (independent variable):** SW state discriminability, Cohen’s *d* of fractional occupancy between WAKE and NREM epochs, z-scored across all sessions
- **M (mediator):** Behavioral arousal, derived as described below
- **Y (dependent variable):** Neural state structure, the first principal component (PC1) of three neural state features (see below), z-scored across all sessions

#### Behavioral Arousal (M)

Behavioral arousal was represented by a composite measure derived from median pupil diameter and median orofacial motion energy, calculated for each HMM state within each session. Each measure was first z-scored within session and then standardized across the pooled observations from all sessions and animals before principal component analysis (PCA). We retained the first principal component (PC1) as the behavioral-arousal score; PC1 explained 84% of the variance across the two behavioral measures. Feature loadings and variance explained by PC1 and PC2 are shown in **Fig. S7A.**

#### Neural State Structure (Y)

Neural state structure was characterized using three HMM state-level features: mean lifetime, Bhattacharyya distance and activity entropy. These features were z-scored and submitted to PCA. The first principal component (PC1, explaining 52% of variance) was retained as the Neural State Structure (Y) and z-scored prior to analysis. Feature loadings on PC1 and PC2 are reported in **Fig. S7B**.

#### Mediation Model

The mediation model comprised three linear mixed-effects regression paths, each fitted using restricted maximum likelihood (REML = False) with animal identity as a random intercept grouping factor, implemented via the *mixedlm* function in the Python *statsmodels* package:

- **Path a (X → M):** Behavioral Arousal regressed on SW State Discriminability
- **Path b (M → Y | X):** Neural State Structure regressed on Behavioral Arousal, controlling for SW State Discriminability
- **Direct effect c′ (X → Y | M):** Neural State Structure regressed on SW State Discriminability, controlling for Behavioral Arousal

The total effect c (X → Y) was estimated by regressing neural state structure on SW state discriminability alone, without controlling for arousal (**Fig. S7E**).

#### Bootstrap Inference

The point estimates and p-values for all path coefficients (a, b, c, c′) reported in the text are derived from the mixed-effects models described above. Statistical significance of the indirect effect was assessed using a non-parametric bootstrap procedure (n = 5,000 iterations) with animal-level resampling to preserve the clustered data structure. On each iteration, animals were resampled with replacement and path coefficients a and b were estimated using ordinary least squares (OLS) regression. The indirect effect a × b was computed for each bootstrap sample (**Fig. S7F**). A 95% percentile confidence interval (CI) was derived from the bootstrap distribution, and a two-tailed p-value was computed as twice the proportion of bootstrap samples on the opposite side of zero from the point estimate. Significance was inferred when the 95% CI excluded zero. A fixed random seed (seed = 42) was used for reproducibility.

### Triplet-based identification of arousal-tuned ensembles

The micro-state triplet framework was defined independently in each session. For each session, the low-motion, high pupil (LM-HP) state was designated as the reference state. Valid micro-state triplets were defined as (lower-arousal state, LM-HP, higher-arousal state) combinations of micro-states for which both the LM-HP → higher-arousal and LM-HP → lower-arousal transition probabilities exceeded chance level, defined as 0.5 /n_states (where n_states is the number of HMM states in that session; **Fig. S9A**). This criterion ensured that only state transitions occurring with above-chance consistency were included in the analysis.

### Activity difference computation

For each valid micro-state triplet, the mean ΔF/F activity of every neuron was computed separately for the reference, higher-arousal, and lower-arousal states. Two activity differences were derived per neuron: Δ_high_ = mean(higher-arousal ΔF/F) − mean(ref ΔF/F) and Δ_low_ = mean(lower-arousal ΔF/F) − mean(ref ΔF/F). Plotting Δ_high_ against Δ_low_ for all neurons placed each neuron in one of four quadrants (Q1-Q4) defined by the sign of each difference (**Fig. 5C-D**).

### Triplet group classification

Triplets were classified using a two-stage strategy combining a permutation-based first-stage criterion with a Pearson correlation consistency check.

In the first stage, a session-level permutation null distribution was generated by randomly shuffling the PM-state assignment of each frame 500 times while preserving the approximate frame counts of each state. For each shuffle, the percentage of neurons falling in each quadrant was recorded, and the 95th percentile across shuffles was used as the significance threshold for that quadrant (see **Fig. S9B** for an example of the observed quadrant percentages relative to this threshold). A triplet passed the first-stage criterion as an opposing triplet if Q2 or Q4 exceeded its 95th-percentile null and (Q2+Q4) > (Q1+Q3); it passed the first-stage criterion as a co-modulated triplet if Q1 or Q3 exceeded its 95th-percentile null and (Q1+Q3) > (Q2+Q4).

In the second stage, the Pearson correlation (R) between Δ_low_ and Δ_high_ across all neurons in the triplet was computed to confirm the population-level geometry. Opposing triplet classification additionally required R < 0 with p < 0.05 (anti-diagonal tilt, confirming opposing tuning); co-modulated triplet classification additionally required R > 0 with p < 0.05 (main-diagonal tilt, confirming co-directional tuning).

A triplet was classified as opposing or co-modulated only if both stages were satisfied; triplets satisfying the first-stage criterion but failing the consistency check (or vice versa) were left unclassified, as were triplets satisfying neither criterion. Opposing and co-modulated triplets were mutually exclusive by construction.

### Ensemble analysis

Within each opposing triplet, neurons were considered significant Q2 or Q4 members if both Δ_high_ and Δ_low_ were significantly different from 0 (Welch t-test, α = 0.01, two-tailed, Welch– Satterthwaite degrees of freedom) and their signs placed them in the respective quadrant. Arousal+ and Arousal− ensembles were then defined by intersecting the significant Q2 and Q4 neuron sets, respectively, across all opposing triplets within a session, regardless of which specific higher-or lower-arousal state was involved in each transition.

To characterize ensemble activity profiles, the mean ΔF/F (z-scored) of Arousal+ and Arousal− ensemble neurons was computed separately for each PM state, ordered from low to high arousal (LM-LP → LM-IP → LM-HP → IM-HP → HM-HP). Cross-session mean trajectories were obtained by averaging across sessions, with variability expressed as SEM.

To assess spatial organization, pairwise Euclidean distances between neuron centroids (in pixels, as recorded in the two-photon field of view) were computed within and across ensemble classes. Within– and across-class distances were compared across sessions using paired t-tests.

### Use of generative AI

During preparation of this manuscript, the authors used ChatGPT (Azure OpenAI, gpt-5.6-terra) and Claude (Anthropic, claude-sonnet-4.6) to assist with language refinement, manuscript organization, and generation of alternative wording for sections of the text. All AI-assisted content was critically reviewed, revised where necessary, and verified by the authors, who take full responsibility for the accuracy, integrity, and final content of the manuscript.

### Data and code availability

All behavioral and neural data related to this study will be deposited in Figshare and made publicly available upon publication. The code used for data processing, analysis, and figure generation will be made available through GitHub and archived in Zenodo upon publication.

## Acknowledgements

We thank Clint Ederveen Janssen and Tjeerd Weijers for assistance in constructing the head-fixed mouse recording setup; Kasper van Nieuwland for help with setting up and aligning the face camera; Réka Kazella, Nikita Cijsouw, María de la Rosa Sanz and Plato Derksen for assistance with mouse habituation; and Mingcan Yu for valuable discussions and suggestions regarding the use of logistic regression for the pupil-based state classifier. We also thank members of the Cognitive and Systems Neuroscience group for helpful discussions and feedback throughout this project.

## Author contributions

Conceptualization: J.Y.W., U.O.; Data curation: J.Y.W.; Formal analysis: J.Y.W.; Funding acquisition: J.Y.W., U.O., C.M.A.P; Investigation: J.Y.W., K.Tijsseling; Methodology: J.Y.W., U.O., G.J.H., K.Takahashi, K.Tijsseling, M.B., A.K.; Software: J.Y.W.; Supervision: U.O., C.M.A.P.; Validation: J.Y.W.; Visualization: J.Y.W.; Writing – original draft: J.Y.W.; Writing – review & editing: J.Y.W., U.O. and all other authors.

## Extended Data

**Figure S1.**
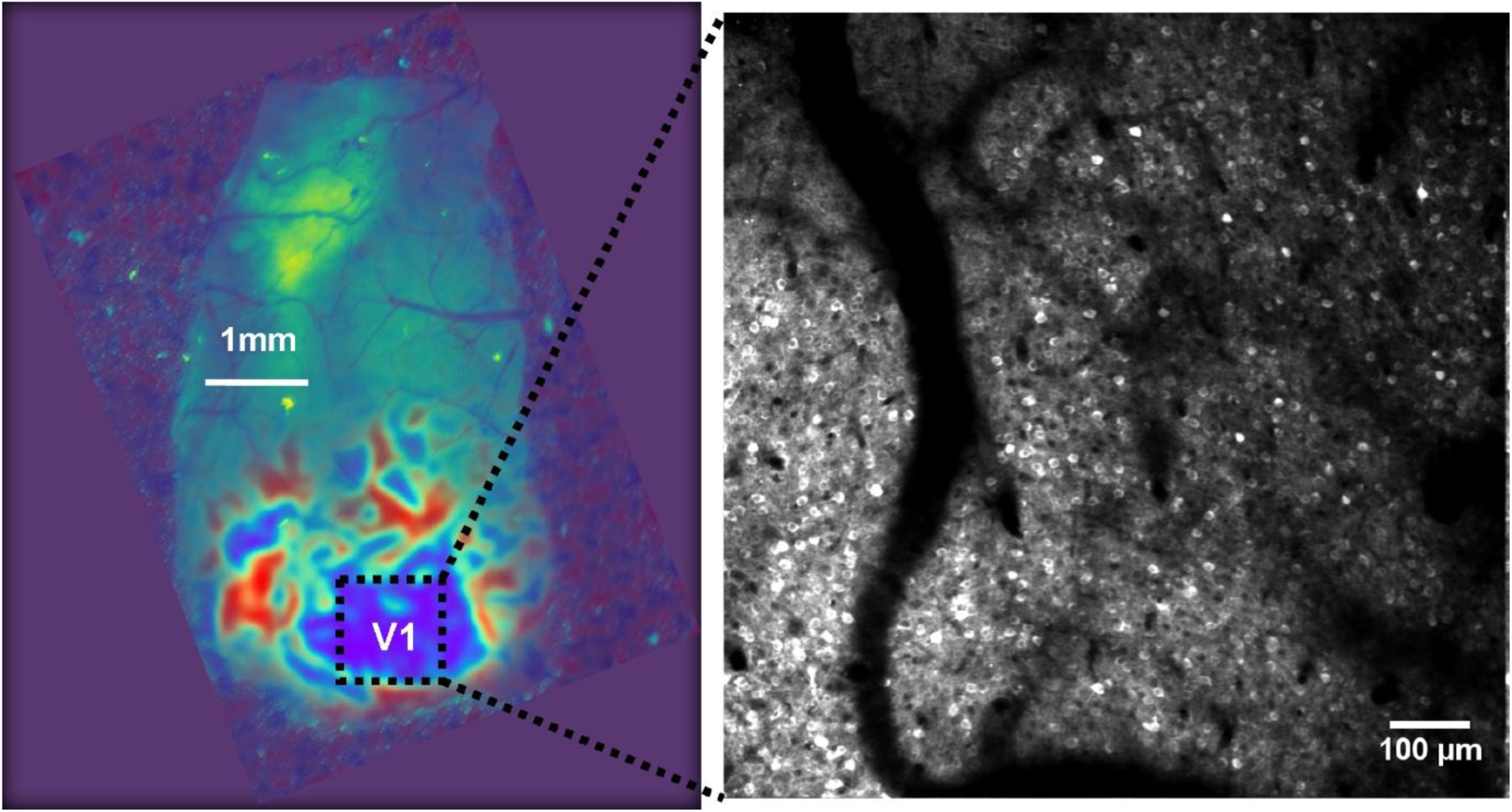
Recording location in primary visual cortex (V1). **Left:** Representative widefield retinotopic sign map used to identify V1. White bar, 1 mm. The white scale bar indicates 1 mm, and the dashed box marks the two-photon imaging field of view (FOV) shown on the right. **Right:** Representative two-photon image of GCaMP6f-expressing layer II/III intratelencephalic (IT) pyramidal neurons in V1. Scale bar, 100 μm.

**Figure S2.**
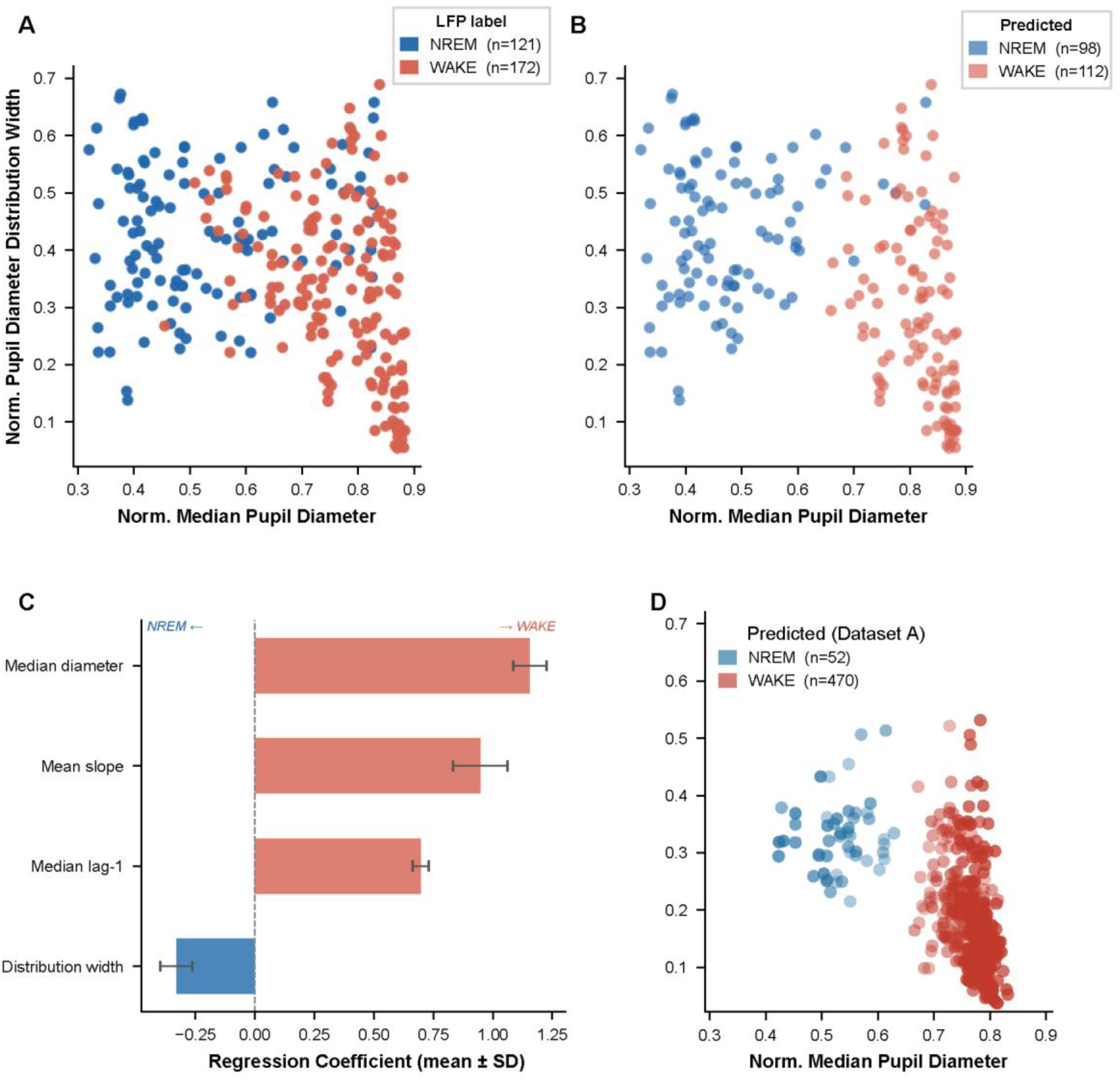
Validation of the logistic regression classifier for pupil-based brain state inference. **(A)** Pupil feature space for a representative session from Dataset B, colored by LFP ground truth label. Each point represents one 60-second epoch. **(B)** Same session and same feature space as (A). Labels are now derived from the logistic regression classifier rather than LFP ground truth. Only high-confidence predictions (P(NREM) ≥ 0.72 or P(WAKE) ≥ 0.72) are shown; epochs below the confidence threshold are excluded. **(C)** Logistic regression coefficients (mean ± SD across cross-validation folds). Median pupil diameter (β = 1.16 ± 0.07), mean slope (β = 0.95 ± 0.12), and lag-1 median (β = 0.70 ± 0.03) were positively associated with WAKE, while distribution width was positively associated with NREM (β = –0.33 ± 0.07). **(D)** Pupil feature space for a representative session from Dataset A (no LFP ground truth available), colored by classifier prediction. Only high-confidence epochs are shown. Point transparency reflects prediction confidence (higher opacity = higher posterior probability).

**Figure S3.**
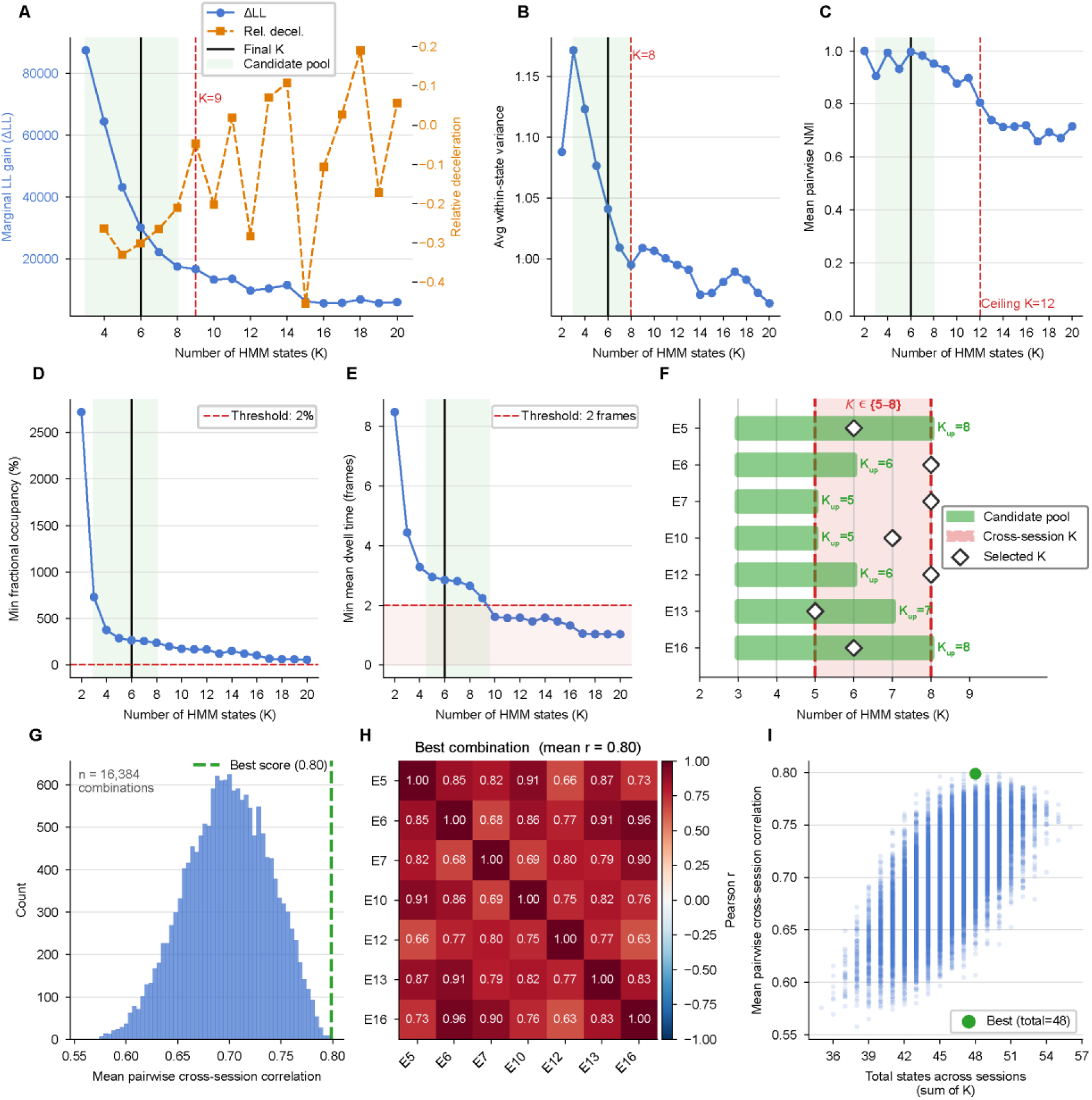
Multi-stage selection of the optimal number of HMM states (K). **(A–E) HMM fitting for a representative session (session E5). (A)** Marginal log-likelihood gain (ΔLL, blue, left axis) and relative deceleration of LL gain (orange, right axis) across increasing numbers of HMM states (K). The red dashed line indicates the LL elbow (K=9). The green shaded area indicates the candidate pool of K values and black vertical line indicates the final selected K (same for B, C, D, E). **(B)** Average within-state variance across increasing numbers of K. The red dashed line indicates the variance elbow (K=8). **(C)** Mean pairwise normalized mutual information (NMI) across repeated model fits. The red dashed line indicates the stability ceiling (K=12), above which additional states yielded limited improvements in model stability. **(D)** Minimum fractional occupancy across states. The red dashed line denotes the occupancy threshold (2%), below which states were considered insufficiently represented. **(E)** Minimum mean dwell time across states. The red dashed line denotes the dwell-time threshold (2 frames). **(F)** Cross-session candidate K ranges. For each imaging session, the candidate pool was defined as the range of K values satisfying all model-quality and biological constraints. The cross-session consensus range (red shaded region, K = 5–8) spans from the most restrictive upper bound across session candidate pools (K_up_ = 5; sessions E7 and E10) to the most permissive upper bound (K_up_ = 8; sessions E5 and E16), representing the range of K values with cross-session support. Diamond markers indicate the final K selected per session following cross-session optimization. **(G–I) Cross-session optimization of K combinations. (G)** Distribution of mean pairwise cross-session correlations across all evaluated K combinations. The green dashed line indicates the best-scoring combination. **(H)** Pairwise cross-session correlation matrix for the optimal K combination. Values indicate Pearson correlation coefficients between Hungarian-matched state feature profiles across sessions**. (I)** Relationship between total model complexity (sum of K across sessions) and mean pairwise cross-session correlation. The green marker indicates the optimal solution identified by the search procedure.

**Figure S4.**
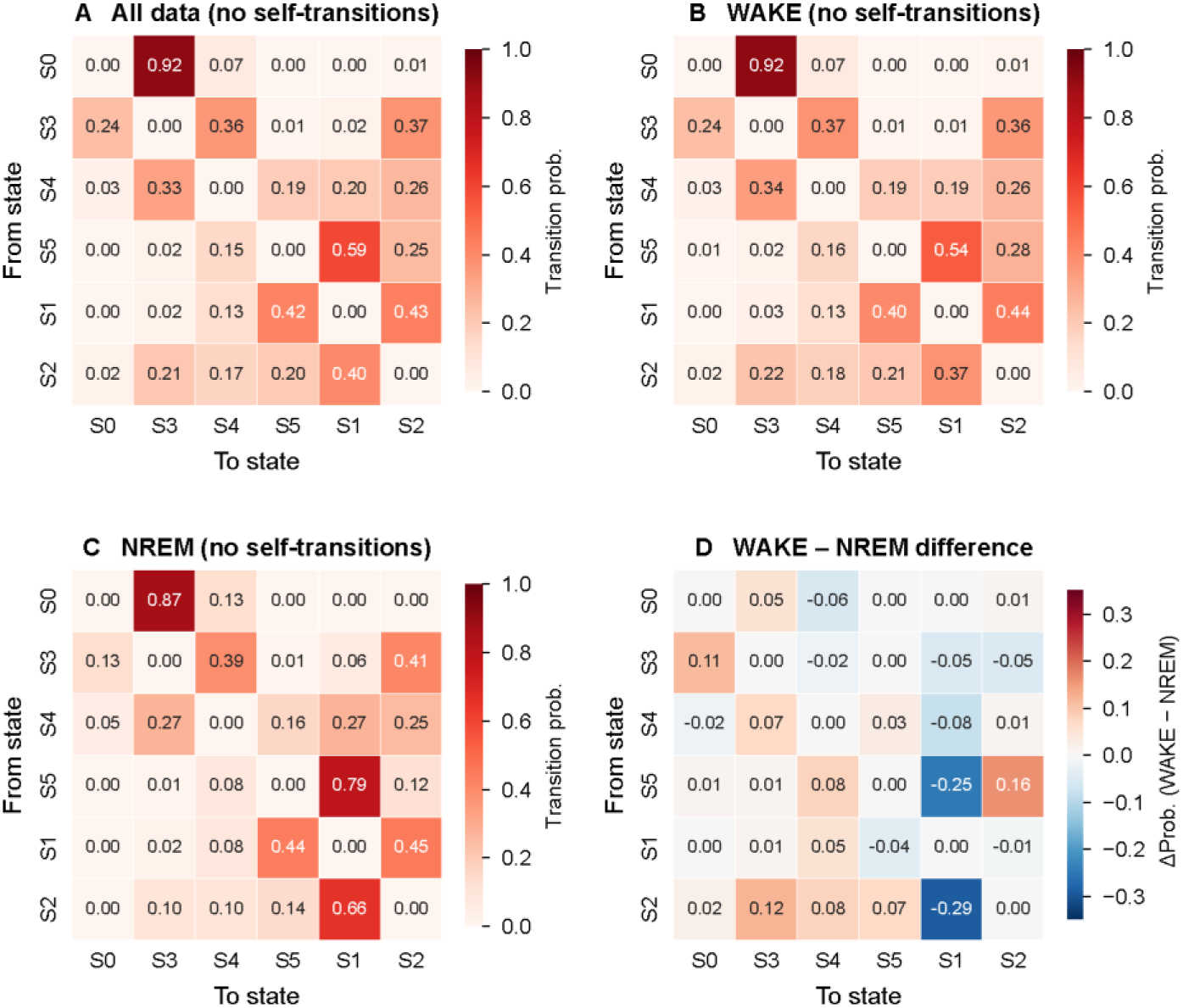
HMM state transition probabilities across WAKE and NREM epochs. **(A)** Transition probability matrix for the representative session (E5; same session as Fig. 3B), computed across all data with self-transitions excluded. **(B, C)** Transition matrices computed separately for WAKE (B) and NREM (C) epochs from the same session (self-transitions excluded). **(D)** WAKE – NREM difference matrix, showing the change in transition probabilities between macro-states (excluding self-transitions). Positive values (red) indicate transitions more probable during WAKE; negative values (blue) indicate transitions more probable during NREM.

**Figure S5.**
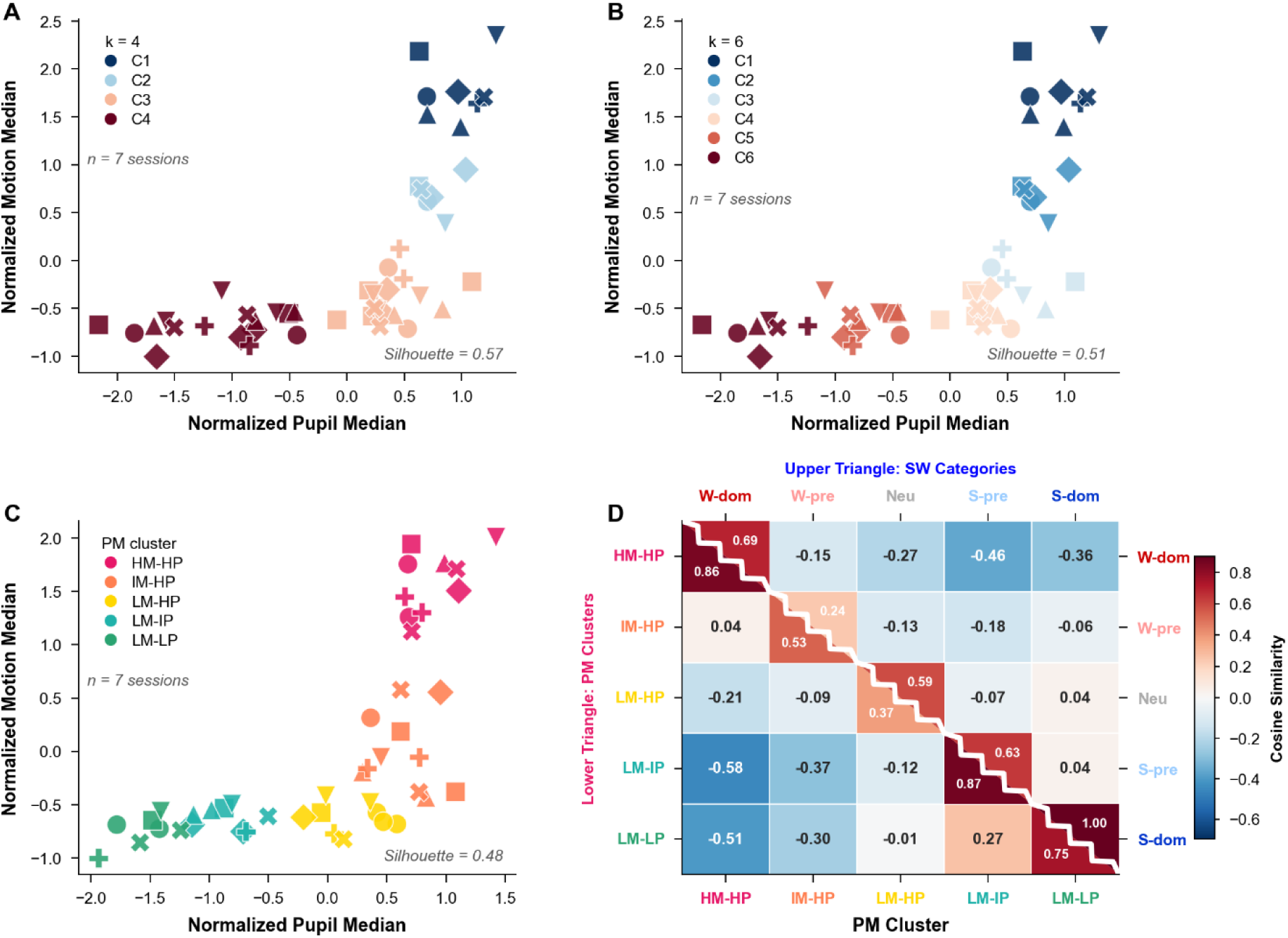
Validation of k=5 as the optimal number of pupil-motion (PM) clusters and robustness of cluster structure across variable HMM state counts. **(A–B)** Pupil-motion feature space with k=4 (A) and k=6 (B) k-means solutions, applied to the same per-state pupil and motion medians used in Figure 4A (n = 7 sessions). Each point represents one HMM state from one session; marker shape denotes animal identity. Silhouette scores are annotated in each panel (k=4: Silhouette = 0.57; k=6: Silhouette = 0.51). **(C)** Pupil-motion feature space using the maximum valid number of HMM states per session, with k=5 clustering applied to the resulting per-state features (n = 7 sessions). Color coding follows the same five semantic PM clusters as Figure 4A. Silhouette score = 0.48. **(D)** Composite cosine similarity heatmap computed from the maximum-state HMM data, following the same split-diagonal format as Figure 4C.

**Figure S6.**
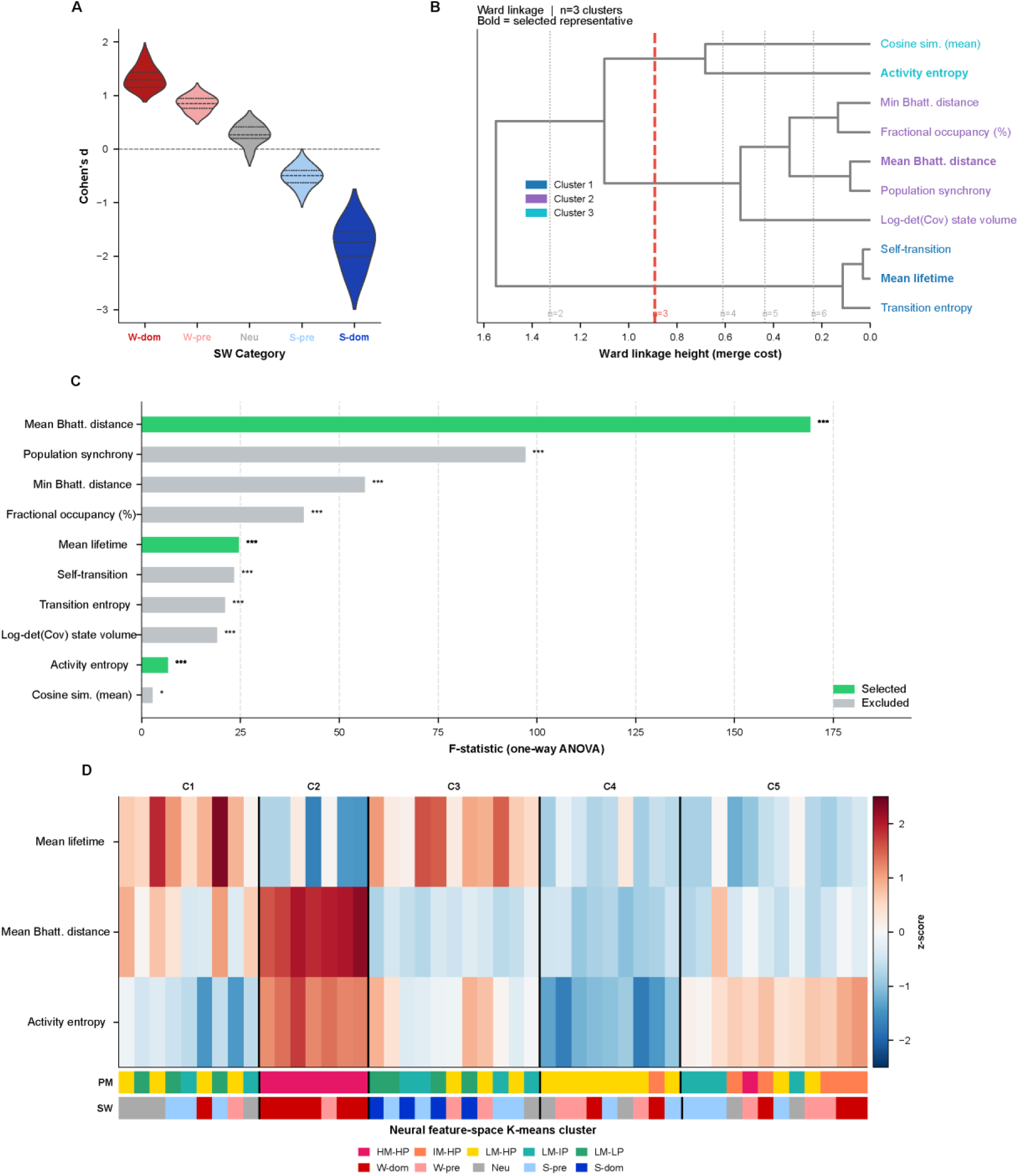
Neural state feature-space characterization and cross-session clustering. **(A)** SW category assignment. Cohen’s d of fractional occupancy between WAKE and NREM epochs was computed for each micro-state and used to assign states to five SW categories via k-means (K = 5): Wake-dominant (W-dom), Wake-prefer (W-pre), Neutral (Neu), Sleep-prefer (S-pre), and Sleep-dominant (S-dom). Violin plots show the distribution of Cohen’s d per category across sessions; dashed line indicates d = 0. **(B)** Hierarchical clustering dendrogram of ten candidate micro-state features using Ward linkage. Three feature clusters were identified by visual inspection (red dashed line); dotted grey lines show alternative cuts (n = 2–6) for reference. Bold leaf labels indicate the representative feature selected from each cluster based on the highest ANOVA F-statistic. **(C)** One-way ANOVA F-statistic for each candidate feature, computed across preliminary K-means clusters. Green bars indicate the three selected representatives (one per feature cluster); grey bars indicate excluded features. Significance levels: * p < 0.05, ** p < 0.01, *** p < 0.001. **(D)** Cross-session K-means clustering of HMM micro-states (K=5) based on the three selected features (panel C, green). Heatmap shows z-scored feature values for all pooled HMM states (n=48 states across 7 sessions), sorted by cluster assignment (C1–C5, separated by vertical black lines). Each column represents one neural state from one session; each row represents one feature. Two annotation bars below the heatmap indicate PM cluster assignment (upper bar) and SW category assignment (lower bar).

**Figure S7.**
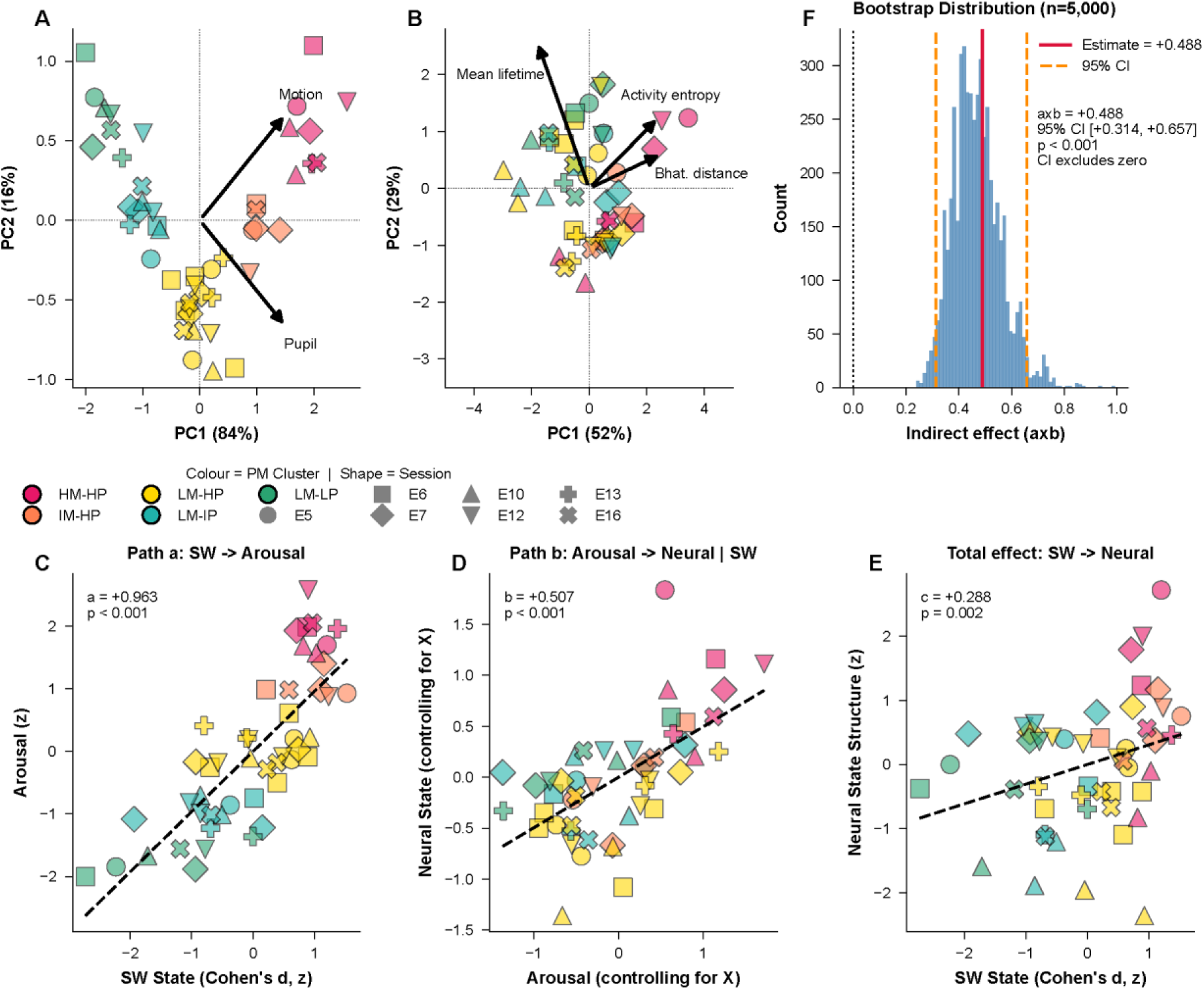
Mediation analysis: variable construction and path estimates. **(A)** Principal component analysis (PCA) of median pupil diameter and median orofacial motion energy used to define the behavioral arousal variable. Data points are projected onto PC1 and PC2 derived from median pupil diameter and median orofacial motion. Arrows indicate feature loadings scaled by explained variance; arrow direction and length reflect the contribution of each feature to each principal component. PC1 explains 84% of variance; PC2 explains 16%. **(B)** PCA scatter plot of neural state features. Data points are projected onto PC1 and PC2 derived from three neural state features: mean lifetime, mean Bhattacharyya distance, and activity entropy. Arrows indicate feature loadings scaled by explained variance. PC1 explains 52% of variance; PC2 explains 29%. **(C)** Scatter plot of SW state discriminability (Cohen’s d, z-scored; X) against the behavioral arousal (z-scored; M), corresponding to path a (X → M) of the mediation model. Dashed line indicates the linear regression fit. Mixed-effects coefficient a = +0.963, p < 0.001. **(D)** Partial residual plot for path b of the mediation model, showing the relationship between the behavioral arousal (M) and Neural State Structure (Y), after partialling out the effect of SW state discriminability (X) from both variables. Dashed line indicates the linear regression fit. Mixed-effects coefficient b = +0.507, p < 0.001. **(E)** Total association between SW state discriminability (X) and neural state structure (Y), corresponding to the total effect c. The dashed line indicates the fitted trend. The annotated coefficient and p-value were obtained from the mixed-effects model with animal identity included as a random intercept. **(F)** Bootstrap distribution of the indirect effect (a×b) based on 5,000 bootstrap samples with animal-level resampling. The red vertical line indicates the point estimate; orange dashed lines indicate the 95% bootstrap confidence interval. Indirect effect a×b = +0.488, 95% CI [+0.314, +0.657], p < 0.001. In all panels, point color indicates PM cluster and point shape indicates animal identity (as in Figure 4A).

**Figure S8.**
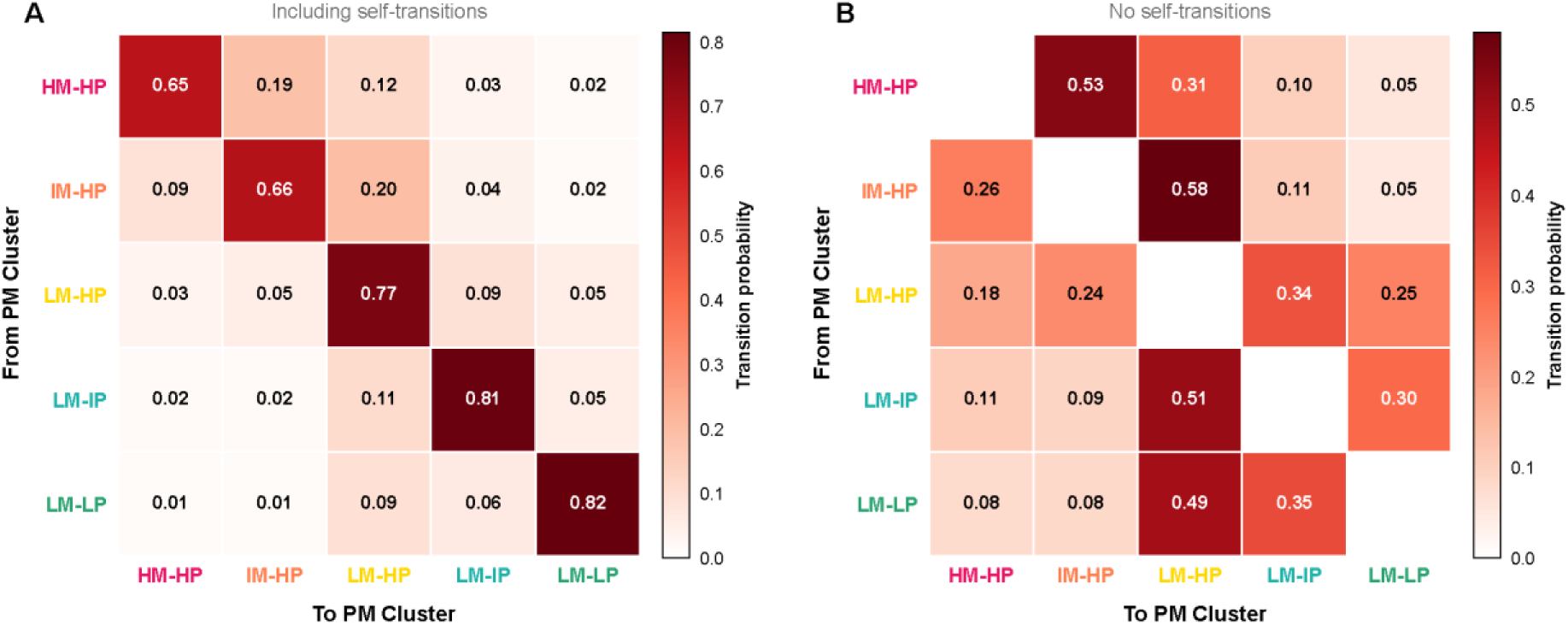
Full transition matrices between PM-derived behavioral clusters. **(A)** Transition probability matrix between PM-derived behavioral clusters, including self-transitions. Each row shows the probability of transitioning from a given PM clusters, and each column shows the probability of transitioning to a given PM cluster. Values indicate transition probabilities averaged across 7 sessions. Clusters are ordered from low to high arousal (LM-LP, LM-IP, LM-HP, IM-HP, HM-HP), as in Figure 5A. **(B)** Same as (A), but with self-transitions excluded and the remaining probabilities renormalized to sum to 1 within each row.

**Figure S9.**
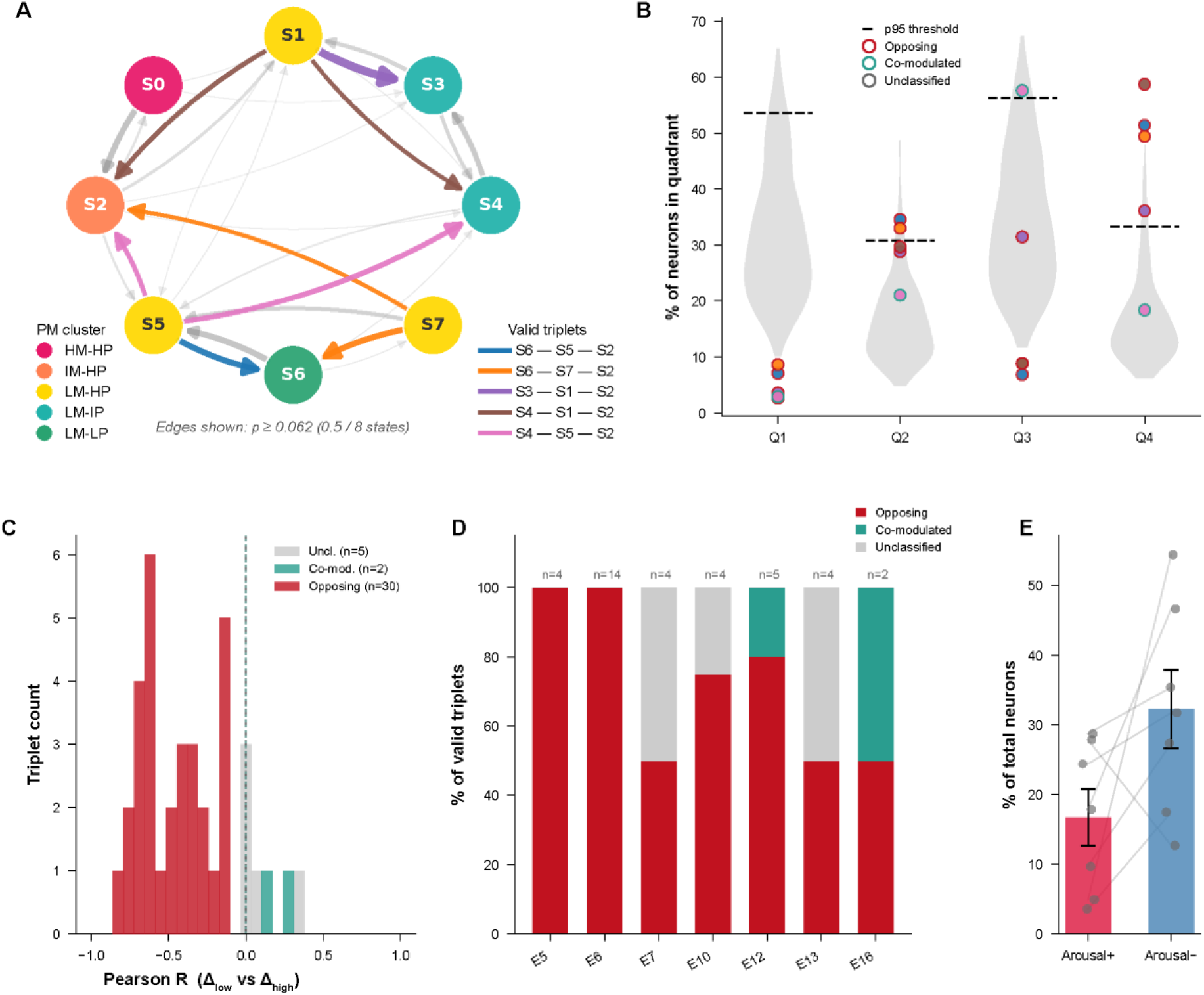
Triplet classification pipeline and identification of arousal-tuned neuronal ensembles. **(A)** Directed transition graph for one example session (E12). Nodes are colored by PM cluster identity (see legend). Only edges with transition probability ≥ 0.062 (0.5 / 8 states) are shown; edge width and opacity are proportional to transition probability. Highlighted edges indicate the LM-HP → higher-arousal and LM-HP → lower-arousal transitions that define each valid triplet; each color identifies one valid triplet (see legend), labelled as lower-arousal state — reference state — higher-arousal state. **(B)** Distribution of the percentage of neurons assigned to each quadrant (Q1–Q4) across all valid state-transition triplets in the same example session (E12). Grey violins show the null distribution of quadrant percentages derived from 500 frame-label shuffles. The dashed horizontal line indicates the 95th percentile of the shuffle-based null, used as the significance threshold for quadrant dominance. Each dot represents one valid triplet; dot fill color identifies the triplet as in (A); dot border color indicates triplet classification: Opposing (red border), Co-modulated (teal border), or Unclassified (grey border). **(C)** Distribution of within-triplet Pearson correlation coefficients between Δ_low_ and Δ_high_, computed across all neurons within each triplet and pooled across sessions. Histogram bars are colored according to triplet group: Opposing (red), Co-modulated (teal) and Unclassified (grey). The vertical dashed line marks zero correlation. **(D)** Fraction of valid triplets classified as Opposing, Co-modulated, or Unclassified, shown separately for each session. Numbers above each bar report the total number of valid triplets (n) in that session. **(E)** Proportion of neurons classified as Arousal+ or Arousal− ensembles across sessions. Bars show mean ± SEM across sessions (n = 7); grey dots represent individual sessions; grey lines connect values from the same session. Arousal+ ensembles: 16.7 ± 4.1%; Arousal− ensembles: 32.2 ± 5.6% of all recorded neurons.

**Figure S10.**
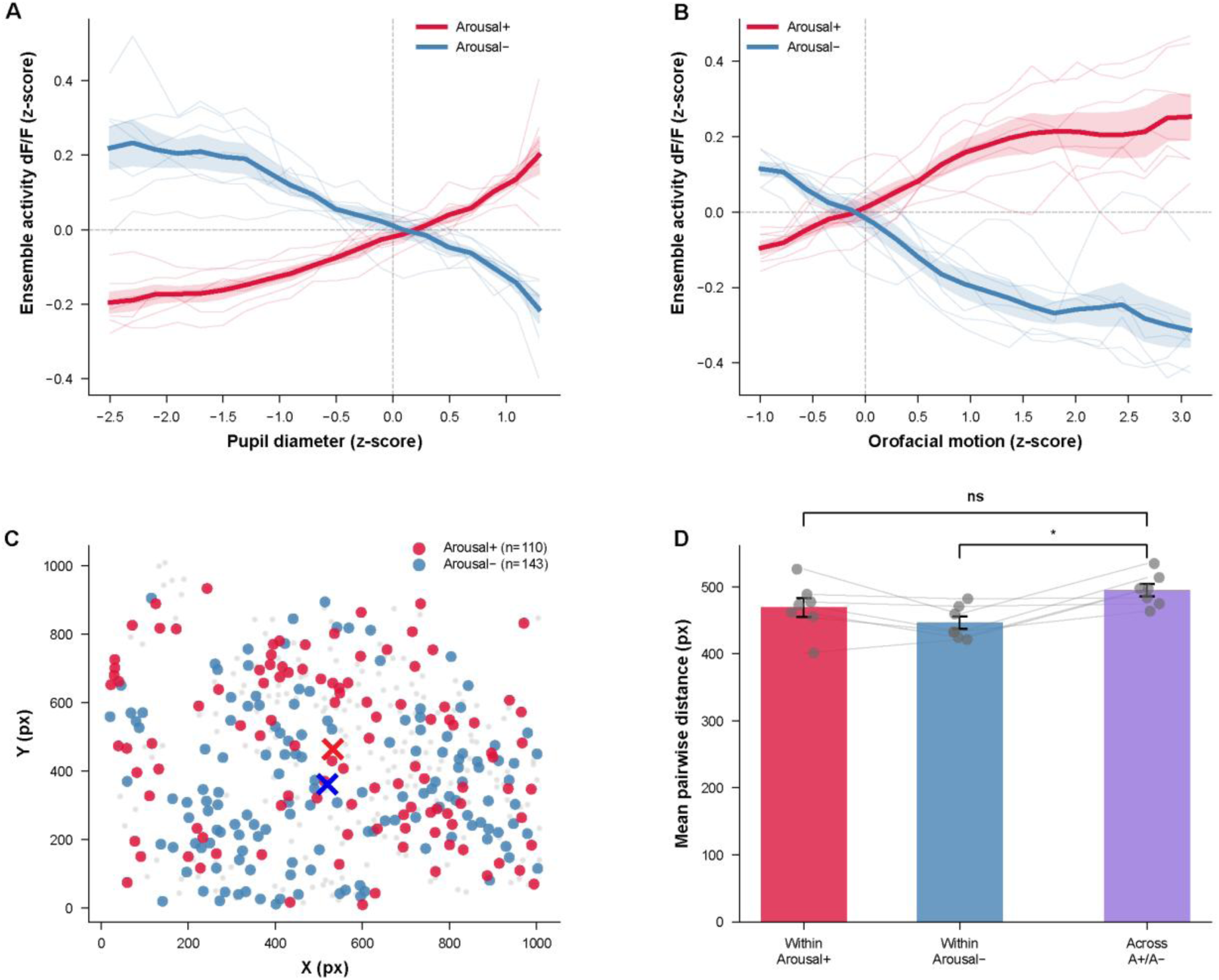
Continuous arousal tuning and spatial organization of Arousal+ and Arousal− ensembles. **(A)** Activity of Arousal+ (crimson) and Arousal− (blue) ensembles as a function of pupil diameter across all imaging frames. Traces show z-scored dF/F averaged across ensemble neurons within each session and then across sessions; thick lines indicate the session average and shaded bands the SEM across 7 sessions, while thin lines show individual sessions. **(B)** Same as (A), but for orofacial motion. **(C)** Spatial distribution of Arousal+ (red) and Arousal− (blue) ensembles within an example field of view. Grey dots denote all other recorded neurons. Crosses indicate the centroid of each ensemble. **(D)** Mean pairwise distances between ensemble neuron locations. Bars show the mean ± SEM across sessions of the mean pairwise distance (in pixels) within Arousal+ ensembles, within Arousal− ensembles, and across Arousal+–Arousal− neuron pairs. Grey dots represent individual sessions; grey lines connect values from the same session. Within– and between-ensemble distances were compared using paired t-tests (ns, not significant; * p < 0.05).

